# Thiamine metabolism genes in diatoms are not regulated by thiamine despite the presence of predicted riboswitches

**DOI:** 10.1101/2022.01.04.474978

**Authors:** Marcel Llavero Pasquina, Katrin Geisler, Andre Holzer, Payam Mehrshahi, Gonzalo I Mendoza-Ochoa, Shelby Newsad, Matthew P Davey, Alison G. Smith

## Abstract

- Thiamine pyrophosphate (TPP), an essential co-factor for all species, is biosynthesised through a metabolically expensive pathway regulated by TPP riboswitches in bacteria, fungi, plants and green algae. Diatoms are microalgae responsible for approximately 20% of global primary production. They have been predicted to contain TPP aptamers in the 3’UTR of some thiamine metabolism-related genes, but little is known about their function and regulation.
- We used bioinformatics, antimetabolite growth assays, RT-qPCR, targeted mutagenesis and reporter constructs to test whether the predicted TPP riboswitches respond to thiamine supplementation in diatoms. Gene editing was used to investigate the functions of the genes with associated TPP riboswitches in *Phaeodactylum tricornutum*.
- We found that thiamine-related genes with putative TPP aptamers are not responsive to thiamine or its precursor 4-amino-5-hydroxymethyl-2-methylpyrimidine (HMP), and the targeted mutation of the TPP aptamer in the HMP-P synthase (*THIC*) does not deregulate thiamine biosynthesis in *P. tricornutum*. Through genome editing we established that *PtSSSP* is necessary for thiamine uptake and that *PtTHIC* is essential for thiamine biosynthesis.
- Our results highlight the importance of experimentally testing bioinformatic aptamer predictions and provide new insights into the thiamine metabolism shaping the structure of marine microbial communities with global biogeochemical importance.

## Introduction

Thiamine pyrophosphate (TPP), the biologically active form of thiamine (vitamin B_1_), acts as a co-factor for key enzymes such as pyruvate dehydrogenase, transketolase and pyruvate decarboxylase and is an essential micronutrient for virtually all organisms (Hanson *et al*., 2018). The widespread use of thiamine across all kingdoms of life suggests a long evolutionary history and supports the hypothesis that B vitamins are remnants of the first organic catalysts in the RNA world (White, 1976). TPP is biosynthesised *de novo* via the condensation of two intermediates: 4-amino-5-hydroxymethyl-2-methylpyrimidine pyrophosphate (HMP-PP) and 4-methyl-5-(2-phosphooxyethyl)thiazole (HET-P), but the production of these precursors follows alternative routes in different kingdoms (Webb *et al*. 2007). In prokaryotes, plants and green algae, the pyrimidine moiety is produced from 5-aminoimidazole ribotide (AIR) by HMP-P synthase (THIC), whereas in fungi it is produced from pyridoxal-5-phosphate (PLP) and histidine by THI5/NMT1 (Coquille *et al*., 2012). HET-P is produced in eubacteria via THIG from iminoglycine, pyruvate, glyceraldehyde-3-phosphate and cysteine, and in archaea, fungi, plants and green algae it is produced via THI4 using NAD^+^, glycine and a sulphur atom from a cysteine residue in the active site (Jurgenson *et al*., 2009). THI4 is thus a suicide enzyme only capable of a single turnover, which is also true for THI5/NMT1. Moreover, THIC has a very low turnover rate and is inhibited by the 5’-deoxyadenosine radical intermediate (Palmer & Downs, 2013), making thiamine biosynthesis a metabolically expensive process (Hanson *et al*., 2018).

Many microbial species, including bacteria and algae, have lost the ability to produce thiamine *de novo*, thus reducing metabolic costs. But in return this renders them dependent on an environmental source of the vitamin or one or more of its precursors (Croft *et al*., 2006). Within the algal lineages, thiamine auxotrophy has evolved multiple times (Helliwell *et al*., 2013) and is widespread in bloom-forming algae, including the picoeukaryotic prasinophytes such as *Ostreococcus tauri* and dinoflagellates. It has been hypothesised that environmental levels of thiamine and its intermediates shape the behaviour of algal blooms and determine microbial community structure with significant implications for oceanic ecosystems and global biogeochemical cycles (Bertrand & Allen, 2012; Gutowska *et al*., 2017). Thiamine auxotrophy is less common in diatoms, a major group of marine microalgae responsible for up to 20% of global primary productivity (Field *et al*., 1998; Rousseaux & Gregg, 2014). Interestingly, these organisms are thought to produce HMP-PP via THIC through a pathway homologous to plants and green algae, but HET-P through the bacterial pathway reliant on THIG (Bertrand & Allen, 2012).

The high metabolic cost of thiamine biosynthesis might also explain the presence of feedback regulation mechanisms in species with a complete biosynthetic pathway. We have previously demonstrated that in the presence of exogenous thiamine, the green alga *Chlamydomonas reinhardtii* downregulates the expression of *THIC* and *THI4* via TPP riboswitches, regulatory elements in mRNA that upon direct binding of a ligand, in this case TPP, trigger a change in genetic expression (Croft *et al*., 2007; Moulin *et al*., 2013). Similarly, riboswitches control *THIC* in plants (Wachter *et al*., 2007), and *THI5/NMT1* and *THIA* (equivalent to *THI1/THI4*) in fungi (Cheah *et al*., 2007). Riboswitches contain two functional units: the aptamer and the expression platform (Roth & Breaker, 2009). The aptamer binds a given ligand with high specificity and often with equilibrium dissociation constants (KDs) in the nanomolar range. Upon binding the substrate, the aptamer undergoes a conformational change that is transduced by the associated expression platform into a change of gene expression. In bacteria, where riboswitches responsive to a range of metabolite ligands are widespread, the expression platform mechanism can involve masking the ribosome binding site, the start codon or termination elements. In eukaryotes, all examples characterised to date are those that respond to TPP, in a mechanism involving alternative splicing (Nguyen *et al*., 2016).

In the past decade, several bioinformatic approaches have been developed to identify putative riboswitches based on sequence information. For instance, Croft *et al*. (2007) analysed sequence conservation between the non-coding regions of the *THIC* gene in the diatoms *Phaeodactylum tricornutum* and *Thalassiosira pseudonana* and identified the presence of a putative TPP riboswitch aptamer in the *THIC* 3’ untranslated region (3’UTR). Later, McRose *et al*. (2014) used the conserved functional motif “CUGAGA” as query against transcripts of thiamine-related genes in combination with secondary structure predictions and homology searches to identify putative riboswitches in the genomes of a wide variety of eukaryotic supergroups (alveolates, stramenopiles, rhodophytes, rhizaria, chlorophytes, prasinophytes, cryptophytes and haptophytes). In their study, they identified putative riboswitches for *THIC* and *SSSP*, encoding a predicted thiamine transporter, in diatoms *P. tricornutum, T. pseudonana, Fragilariopsis cylindrus* and *Pseudonitzschia multiseries*.

In diatoms, both the thiamine biosynthetic pathway and the presence of TPP aptamers have been predicted using bioinformatic methods, but they have not been experimentally studied before now. Here, we experimentally test whether the predicted TPP riboswitches found in diatoms *P. tricornutum* and *T. pseudonana* regulate thiamine biosynthesis at the transcript, protein and intracellular thiamine levels using wildtype and reporter strains. In addition, we use a CRISPR/Cas9 approach to test the predicted function of genes containing TPP riboswitches in *P. tricornutum*.

## Materials and Methods

### Strains and culture conditions

*Phaeodactylum tricornutum CCAP 1055/1* was grown in f/2 minus silica without vitamins at 18°C and 30 μmol m^−2^ s^−1^ in a 16:8 hours day-night cycle. *Thalassiosira pseudonana 1085/12* was grown in f/2 plus silica and 0.6 μM cyanocobalamin (Millipore-Sigma) at the same temperature and light regime. *Chlamydomonas reinhardtii UVM4* (Neupert *et al*. 2009) was grown in TAP without vitamins at 24°C and same light regime. Cultures were supplemented with thiamine (Acros Organics, USA), pyrithiamine (Sigma-Aldrich, USA), or 4-amino-5-phosphonooxymethyl-2-methylpyrimidine (HMP)(Sigma-Aldrich, USA), at the indicated concentrations for each of the experiments. Zeocin (InvivoGen, USA) at 75 mg L^−1^ was used to select transgenic *P. tricornutum* cells and at 10 mg L^−1^ to select for *C. reinhardtii* transformants. Cell growth was measured as OD_730_ with a ClarioStar plate reader (BMG Labtech, Germany).

### Prediction of TPP aptamers in newly sequenced diatom genomes

The sequence spanning from the 3’ strand of the P2 stem to the 3’ strand of the P4 stem of eight previously predicted TPP aptamers in diatoms (Croft *et al*., 2007; McRose *et al*., 2014; Table S1a) were used to create hidden Markov models (HMM) for both the forward and the reverse complement. The models were generated by multiple sequence alignments using MAFFT (v7.475; Katoh & Standley, 2013), followed by HMM model construction in HMMER (v3.1b2; Eddy, 2011). Both profiles were searched for against a custom sequence database of diatom genomes (see Table S2) using HMMER’s ‘*hmmsearch*’ function with default parameters. Resulting hits were validated manually and their immediate upstream or downstream open reading frame was annotated through a Pfam search and a reciprocal TBLASTN with *P. tricornutum* (Table S1b). The secondary structure for each predicted aptamer associated with a thiamine-related gene was annotated manually with the assistance of the RNAfold web server tool (Hofacker, 2003; Table S1c). PolyA signal site prediction on *PtTHIC* 3’UTR was performed with the PASPA server using default parameters (Ji *et al*. 2015).

### Identification of thiamine biosynthesis capacity in available diatom genomes

A Benchmarking Universal Single-Copy Ortholog analysis (BUSCO, v5.1.2) was performed using the genome mode to assess completeness of the various assemblies in our custom diatom genome database (Table S2, Seppey *et al*., 2019). Only assemblies with more than 88% (stramenopiles_odb10) and 55.7% (eukaryota_odb10) complete BUSCOs were analysed further. Nucleotide assembly files from the resulting 19 diatom genomes were used to construct BLAST databases. TBLASTN searches (BLAST+, v2.6.0+) were performed for members of all KEGG orthologues associated with thiamine metabolism (KEGG:ko00730) using reference peptide sequences recovered from the KEGG database (Kanehisa & Goto, 2000; Kanehisa, 2019; Kanehisa *et al*., 2021) as queries (Table S3a). In addition, we also performed TBLASTN searches with the predicted thiamine biosynthesis proteins THIC, TH1, THIS, THIO, THIG, THIF, DXS, TPK1, THI4, THIM as well as the thiamine-related proteins SSSP, SSUA/THI5-like and TENA from *P. tricornutum* or *C. reinhardtii* (Table S3c). The best hit for each genome-protein pair was extracted, with full results reported in Table S3b+d. For species with available annotation data, the overlap of TBLASTN results with genomic loci was determined. In those cases where annotation data was lacking or hits did not directly overlap with a genetic locus, contig/scaffold name together with start and end coordinates of the hit are provided. Categorising presence of thiamine biosynthesis genes in a diatom genome was performed based on hits found by either one of the described TBLASTN searches with an E-value cut-off of ≤ 10^−20^, while an E-value between 10^−3^ and 10^−20^ indicated potential existence (Table S3e). Predicted peptide sequences containing NMT1 domains were analysed by multiple sequence alignments in MEGA-X v.10.1.1 (Kumar *et al*., 2018) using MUSCLE (Edgar, 2004) with default parameters and the phylogenetic tree was generated with the default Maximum-Likelihood algorithm and 100 bootstrap iterations.

### RNA Isolation and Protein Isolation

RNA was extracted from liquid nitrogen-frozen cell pellets from 20 mL cultures grown to early stationary phase using the RNeasy Plant Mini Kit (Qiagen, Germany). Immediately after extraction, the RNA samples were treated with 1 U of TURBO DNase (Thermo Fisher Scientific, USA) for 30 minutes before cDNA synthesis. Total protein extracts were obtained from 150 mL cultures grown to early stationary phase by resuspending in X mL of 0.2 M sorbitol (Sigma-Aldrich, USA), 1 % β-mercaptoethanol, and 0.8 M Tris-HCl pH 8.3 (Sigma-Aldrich, USA), where X is equal to the culture OD_750_ before harvesting.

### Analysis of Gene Expression by quantitative PCR

First strand cDNA was generated with SuperScript III reverse transcriptase (Thermo Fisher Scientific, USA) primed with random hexamers. Quantitative PCR was performed with SybrGreen JumpStart Taq (Sigma-Aldrich, USA) in a RotorGene qPCR thermocycler (QIAGEN, Germany) for 40 cycles of 94°C for 20 seconds, 55°C for 20 seconds, and 72°C for 30 seconds (see primers in Table S4). Total transcript levels of genes of interest were normalised to the levels of housekeeping genes histone 4 (H4), ubiquitin conjugating enzyme (UBC) and ubiquitin (UBQ). Relative expression was calculated using the Delta-Delta Ct method adjusted by amplification efficiency. Measurements with amplification efficiency lower than 1.525 (1.67 for cobalamin supplementation experiment) were excluded. Housekeeping genes showing significant differences between treatments were not used for normalisation.

### 3’RACE

First strand cDNA was synthesised using a polyT-VN primer with two anchor nucleotides at its 3’ end and a universal adaptor in its 5’ UTR (see Table S4) (Beilharz & Preiss, 2009). The cDNA was diluted 1/8 in nuclease-free water and used as template for a first touch-down RT-PCR reaction primed with a high-specificity primer (71°C annealing Tm) and a universal reverse primer using Q5 High-Fidelity polymerase (New England Biolabs, USA). The PCR product of this first RT-PCR was then diluted 1/100 in nuclease-free water and used as a template for a semi-nested RT-PCR using a gene-specific primer and the universal reverse primer. Q5 polymerase was used again during 35 cycles using annealing temperature of 65°C and 30 seconds extension. RT-PCR products were run in a 2 % agarose gel at 130 mV for 25 minutes unless otherwise stated. Selected bands were cut, purified with the Illustra^™^ GFX^™^ PCR DNA and Gel Band Purification Kit (Sigma-Aldrich, USA) and sent for Sanger sequencing (Source Bioscience, UK).

### Plasmid construction and algae transformation

All constructs were cloned following the MoClo Golden Gate system (Engler *et al*., 2014). Level 0 parts were reused from existing *P. tricornutum* constructs, from the *C. reinhardtii* MoClo Kit (Crozet *et al*., 2018), or were amplified from *P. tricornutum* genomic DNA using Q5 High Fidelity polymerase (see Table S4 for primers). Level 1 constructs were assembled by *Bsa*I restriction-ligation of Level 0 parts. Level 2 constructs were assembled by *Bpi*I restriction-ligation of Level 1 constructs. Constructs used for CRISPR/Cas9 genome editing were cloned following the sgRNA design strategy described in Hopes *et al*. (2017) and homologous recombination regions were designed to be around 800 bp long and flank the coding sequence of the gene of interest. The level 1 plasmid encoding a Cas9-YFP expression cassette (pICH47742:PtFCP:Cas9YFP), the level 0 plasmid containing the *PtU6* promoter to drive expression of the sgRNAs (pCR8/GW:PtU6) and the plasmid used as template to amplify the sgRNA scaffold (pICH86966∷AtU6p∷sgRNA_PDS) were a kind gift from Dr Amanda Hopes and Prof. Thomas Mock (UEA, UK) and are available on Addgene (Hopes *et al*., 2016).

*C. reinhardtii* transformation was carried out as described in Mehrshahi *et al*. (2020) and *P. tricornutum* transformation as in Yu *et al*. (2021). For co-transformation of plasmids in *P. tricornutum* 2.5 μg of each plasmid was used. For each construct, up to 96 primary zeocin-resistant transformants were initially selected for PCR genotyping and preliminary phenotyping, and then a subset was taken for further characterisation.

### Determination of intracellular vitamin quotas

Cell pellets were harvested from 30 mL cultures 5 days post-inoculation, washed three times with distilled water, and fresh weight of the final pellets was measured before flash freezing in liquid nitrogen and storing at −80°C. Pellets were treated with 250 μL 1 % (v/v) trichloroacetic acid (TCA) (Sigma-Aldrich, USA) and centrifuged at 10,000*g* for 10 minutes recovering the supernatant. TPP and thiamine were then derivatised by mixing 50 μL of the cell extract with 10 μL of freshly prepared 30 mM potassium ferricyanide (Sigma-Aldrich, USA) in 15% (w/v) sodium hydroxide, 15 μL 1 M NaOH, and 25 μL methanol (HPLC-grade; Sigma-Aldrich, USA). The derivatisation mix was centrifuged at 4,000*g* for 10 minutes, and 20 μL of the supernatant were injected for HPLC analysis. An Accela HPLC setup (Thermo Fisher Scientific, USA) was used with a C18 150 x 4.6 mm column (Phenomenex, USA). The fluid phase flowed at 1 mL min^−1^ with a gradient of 5% methanol up to 47.5% at 10 minutes, 100% at 11 minutes, 100% at 15 minutes, 5% at 16 minutes and equilibration at 5% methanol until 21 minutes. The thiamine and TPP derivatives were measured using a Dionex UltiMate 3000 fluorescence detector (Thermo Fisher Scientific, USA) with 375 nm excitation and 450 nm emission. The sensitivity of the fluorescence detector was set at 1 for the first 5 minutes of the HPLC programme and increased to 8 for the rest of the programme. The area of the TPP earlier-half peak at 1.35 minutes and the thiamine peak at 2.1 minutes were used to calculate the amount of each vitamer relative to their respective standard curves.

### Western Blots

Crude protein extracts were mixed with 1% Sodium Dodecyl Sulphate (SDS; Sigma-Aldrich, USA) and boiled for 1 min. The samples were centrifuged at 16,000*g* for 2 minutes, and 15 μL were loaded in a 15% Acrylamide SDS-polyacrylamide gel electrophoresis (SDS-PAGE). The electrophoresis was run at 150 mV for 90 minutes. The proteins were then transferred to a polyvinylidene difluoride (PVDF) membrane applying 20 mA for 20 minutes in a semi-dry transfer cell (Bio-Rad Laboratories, USA). The membrane was blocked in 0.5% powdered milk in TBS-T buffer at 4°C overnight, then incubated for 1 hour with a rabbit anti-HA primary antibody (H6908, Sigma-Aldrich, USA) in 2.5% powdered milk in TBS-T, washed 4 times with TBS-T, then incubated for 1 hour with a goat anti-rabbit secondary antibody conjugated with a Dy800 fluorophore (SA5-35571, Thermo Fisher Scientific, USA) in 2.5% powdered milk in TBS-T. The membrane was finally washed 4 times in TBS-T and once in TBS before being imaged in a fluorescence scanner (Odyssey; Li-Cor Biosciences, USA).

## Results

### Putative TPP aptamers can be found with high conservation in diatom genomes

To analyse the conservation and prevalence of putative TPP aptamers in diatoms, we searched for them in 23 available diatom genomes that were well assembled and annotated (Table S2). We performed HMM searches with a motif based on eight previously predicted diatom TPP aptamer sequences in *P. tricornutum, T. pseudonana, F. cylindrus* and *P. multiseries* (Croft *et al*., 2007; McRose *et al*., 2014; Table S1a). We found a total of 40 new putative TPP aptamers (Table S1b). An additional, more targeted, search for the universally conserved “CUGAGA” motif in the UTRs of annotated *THIC* and *SSSP* genes revealed a putative TPP aptamer in the 3’UTR of *Psammoneis japonica THIC* that had not been detected by the HMM motif search.

All putative diatom TPP aptamers are found in 3’UTRs and share a strong sequence conservation of the P2, P4 and P5 stems as well as a structurally conserved P3a stem of variable length (Fig. **1a**; Table S1c). The P1 stems at the 3’ end of the putative aptamers are generally A-rich and, in *PtTHIC (Phatr3_J38085)*, the P1 stem overlaps with the “AACAAA” motif that has been predicted to be the most likely polyadenylation site in the gene 3’UTR by the PASPA software (Ji *et al*., 2015; Fig. **1b**). The “CUGAGA” motif and overall secondary structure architecture is conserved between diatoms and other aptamers demonstrated to be functional in green algae (Croft *et al*., 2007), plants (Wachter *et al*., 2007) and fungi (Cheah *et al*., 2007) (Fig. **1b**). The P4/5 stem sequence is also well conserved between aptamers from the different groups. In contrast, the P2 stem differs between diatoms and other characterised TPP riboswitches. While in green algae and plants this includes the “AGGG” sequence, which includes the alternative splicing acceptor (AG) used in the mechanism of action determined experimentally (Croft *et al*., 2007; Wachter *et al*., 2007), diatoms have a P2 stem with a conserved “GCGG” sequence, with no obvious AG splicing acceptor nearby in the aptamer.

**Figure 1.**
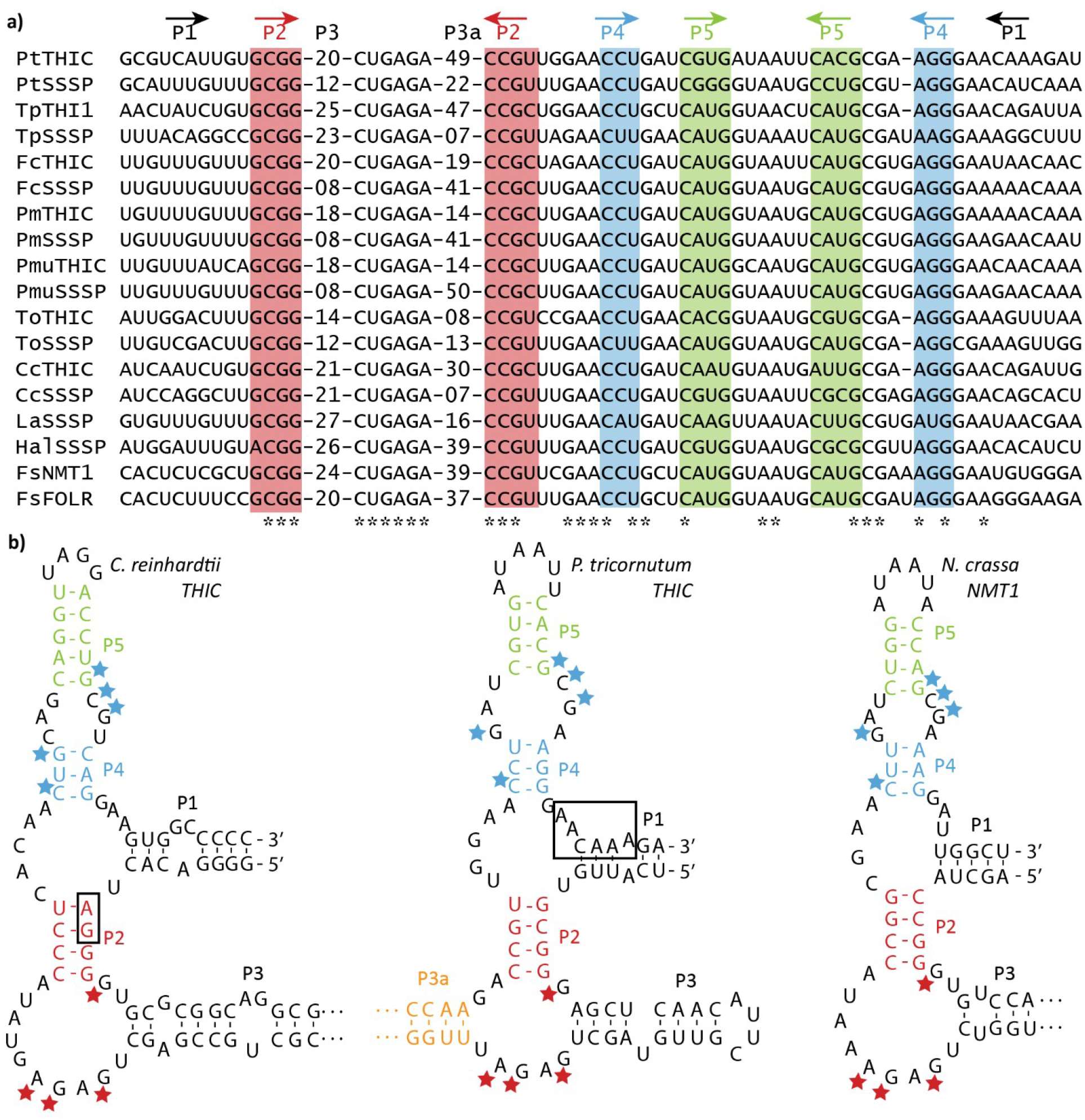
Multiple sequence alignment of 16 predicted diatom thiamine pyrophosphate (TPP) aptamers and structural comparison with previously characterised eukaryotic riboswitches. **(a)** Multiple sequence alignment of previously identified (first eight) and a sample of newly identified TPP aptamers in diatoms. Stems are indicated with arrows and colour coded. See Table S1b and S1c for the full sequences of all predicted TPP aptamers. (b) Structural comparison of the predicted Phaeodactylum tricornutum THIC aptamer (centre) with experimentally described TPP aptamers in Chlamydomonas reinhardtii (left, Croft et al., 2007) and Neurospora crassa (right, Cheah et al., 2007). The pyrimidine-binding residues (“CUGAGA” motif, red stars) and the pyrophosphate-binding residues (“GCG” motif, blue stars) are highlighted. Green algae and plant aptamers contain an alternative 3’ splicing site used in their mechanisms of action in their P2 stem (AG, boxed). The “AACAAA” sequence overlapping with the PtTHIC aptamer P1 stem (boxed) is predicted to be the most likely polyadenylation site by the PASPA software (Ji et al., 2015). *Pt*: Phaeodactylum tricornutum*; Fc*: Fragilariopsis cylindrus*; Tp*: Thalassiosira pseudonana*; To*: Thalassiosira oceanica*; Cc*: Cyclotella cryptica*; Pm*: Pseudonitzschia multiseries; *Pmu*: Pseudonitzschia multistriata*; La*: Licmophora abbreviata*; Hal*: Halamphora sp. MG8b*; Fs*: Fistulifera solaris.

Thirty-one (78%) of the newly predicted aptamers were directly associable with a potential genetic locus involved in thiamine metabolism, predominantly *THIC* and *SSSP*. Overall, TPP aptamers were associated with 11 of 15 identified *THIC* genes and 12 of the 13 identified *SSSP* genes (Fig. **2a**). In addition, putative TPP aptamers were found in genes encoding FOLR domains (folate receptor domain, PF03024) in *Halamphora sp. MG8b* and *F. solaris*. Proteins with FOLR domains in *F. cylindrus, Nitzschia sp. Nitz4* and *Bacillariophyta sp*. (ASM1036717v1), but not in *F. solaris FOLR*, are predicted to have a signal peptide by SignalP 4.1, suggesting a potential role in transport or sensing. Predicted TPP aptamers were also found associated with multiple copies of *F. solaris* genes encoding an NMT1 domain (PF09084; No Message in Thiamine; Maundrell, 1990) (Fig. **2a**; Table S2c). The bioinformatics analysis also provided the means to construct the complete pathway of thiamine metabolism in diatoms indicating both the biosynthetic and salvage routes for provision of the active cofactor, TPP (Fig. **2b**), confirming that synthesis of the pyrimidine moiety uses THIC, as in plants and green algae (as well as bacteria), but that the thiazole group is the bacterial route via ThiG, rather than THI4/THI1 as in all other eukaryotes.

**Figure 2.**
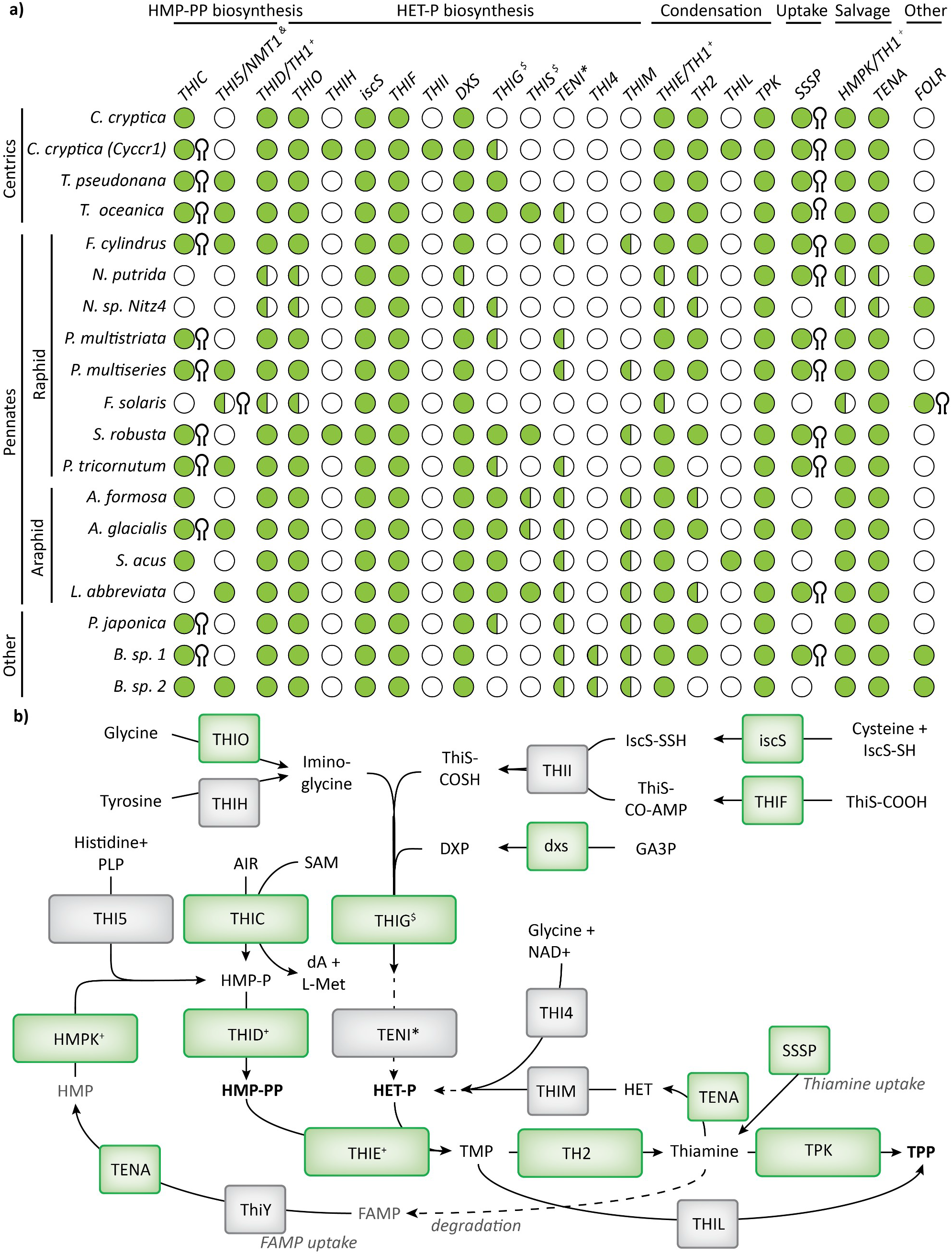
Proposed routes for thiamine biosynthesis in diatoms. **(a)** A TBLASTN search using selected algal peptide sequences as queries (See Table S3c) was performed against 19 diatom genomes to determine the presence (full circle p-value > 10^−20^; half-full circle p-value > 10^−3^) or absence (empty circle) of different thiamine-related genes. The presence of an associated predicted riboswitch in the 3’UTR of the gene is indicated with a hairpin symbol at the right of the circle. The genome abbreviations, accession numbers and references can be found in Table S2. **(b)** Potential thiamine biosynthetic, salvage and uptake routes in diatoms. The pathway steps with strong support across the diatom lineage are shown in green. AIR: 5-Aminoimidazole ribotide; SAM: S-Adenosyl methionine; dA: 5’-deoxyadenosine; L-Met: L-Methionine; GA3P: Glyceraldehyde 3-phosphate; HMP-P: hydroxymethyl-pyrimidine phosphate; HMP-PP: hydroxymethyl-pyrimidine pyrophosphate; HET-P: hydroxyethyl-thiazole phosphate; FAMP: N-formyl-4-amino-5-aminomethyl-2-methylpyrimidine; DXP: 1-deoxy-D-xylulose 5-phosphate; PLP: pyridoxal 5’-phosphate; NAD: nicotinamide adenine dinucleotide; TMP: thiamine monophosphate; TPP: thiamine pyrophosphate. ^&^ THI5/NMT1 candidates contain an NMT1 pfam domain (PF09084). ^$^THIG and THIS are encoded in the chloroplast in P. tricornutum, so the results can be biased in genomes that do not include chloroplast sequences. ^+^THID, THIE and HMPK functions are performed by a single peptide in diatoms (TH1). *In some bacteria TenI accelerates a thiazole tautomerisation reaction, but it is not necessary to synthesise HET-P (Hazra et al., 2011).

In addition to THIC, 9 out of 19 diatom genomes queried revealed at least one gene that encoded a protein with an NMT1 domain. These are associated with THI5, an HMP-P synthase in fungi, but they also have structural homology with ThiY, a bacterial periplasmic component of a pyrimidine precursor ABC transporter (Bale *et al*., 2010). To investigate whether the diatom candidates with NMT1 domains showed closer similarity to THI5 or ThiY, we aligned 10 diatom protein sequences with NMT1 domains with *Bacillus halodurans* ThiY, *S. cerevisiae* THI5, *N. crassa* NMT1 and peptide sequences with NMT1 domains previously identified in other algal species (McRose *et al*., 2014). Multiple sequence alignment and subsequent phylogenetic tree analysis showed that the diatom candidates clustered with the haptophyte (*Emiliana huxleyi*) and cryptophyte (*Guillardia theta*) candidates in a single branch, except for two candidates found in *F. solaris*, which clustered with the chlorophyte peptides (Fig. **S1a**). However, the phylogenetic analysis failed to resolve whether the algal proteins containing NMT1 domains are more closely related to THI5 or ThiY, with bootstrap values all <60. The multiple sequence alignment also revealed that the diatom candidates conserve only 4 of the 15 active site residues in THI5, and 4 out of 8 active site residues in ThiY (Fig. **S1b**). Additionally, except *F. solaris*, all diatom candidates show an extended N-terminus as in ThiY, which is predicted to be a signal peptide by SignalP v.4.1 (Petersen *et al*., 2011). To test whether diatom candidates with NMT1 domains are expressed and regulated by its putative metabolic products, we used RT-qPCR to measure the transcript levels of the *P. tricornutum* and *T. pseudonana* candidates (*Phatr3_J33535* and *THAPS_6708* respectively) in the presence or absence of thiamine or HMP supplementation. The results confirmed the candidates are expressed in both species, but they are not regulated by thiamine (Fig. **S2**). Finally, we used CRISPR/Cas9 to generate *P. tricornutum* mutants with a deletion of the gene coding for an NMT1 domain. Two independent mutants showed no obvious phenotype compared to wild type and could grow in the absence of exogenous thiamine (Fig. **S3**).

### *THIC* transcript levels are unaffected by exogenous thiamine and *P. tricornutum* and *T. pseudonana* are resistant to pyrithiamine

To investigate whether putative riboswitches in other thiamine-related genes in *P. tricornutum* and *T. pseudonana* respond to exogenous thiamine, an RT-qPCR experiment was carried out with cells grown in the presence and absence of 10 μM thiamine or 10 μM HMP, both of which reduce expression of *THIC* in *C. reinhardtii* (Moulin *et al*., 2013). A previous transcriptomics and proteomics study (Bertrand *et al*., 2012) had shown that *PtTHIC* was affected by growth of cells in cobalamin (vitamin B_12_), so this was also included. As expected, in *P. tricornutum PtTHIC* levels dropped about two thirds (p-value 0.03) in the presence of cobalamin relative to the unsupplemented condition (Fig. **3**). The positive control, *PtMETE* (Helliwell *et al*., 2011), showed a 97% reduction (p-value 0.01). In contrast, neither thiamine or HMP supplementation caused significant changes in transcript levels of *PtTHIC* or *PtSSSP (Phatr3_J50012)*. Similarly in *T. pseudonana, TpTHIC (THAPSDRAFT_41733)* transcript levels were unaffected when cells were cultured with 10 μM thiamine. However, in contrast to *P. tricornutum*, thiamine supplementation resulted in approximately one third downregulation (p-value 0.03) of *TpSSSP (THAPSDRAFT_20656)*. To rule out the possibility that these results were explained by the inability of exogenous thiamine to enter the cells, we performed a thiamine uptake test and confirmed that thiamine is actively taken up and metabolised to TPP in *P. tricornutum* (Fig. **S4**).

**Figure 3.**
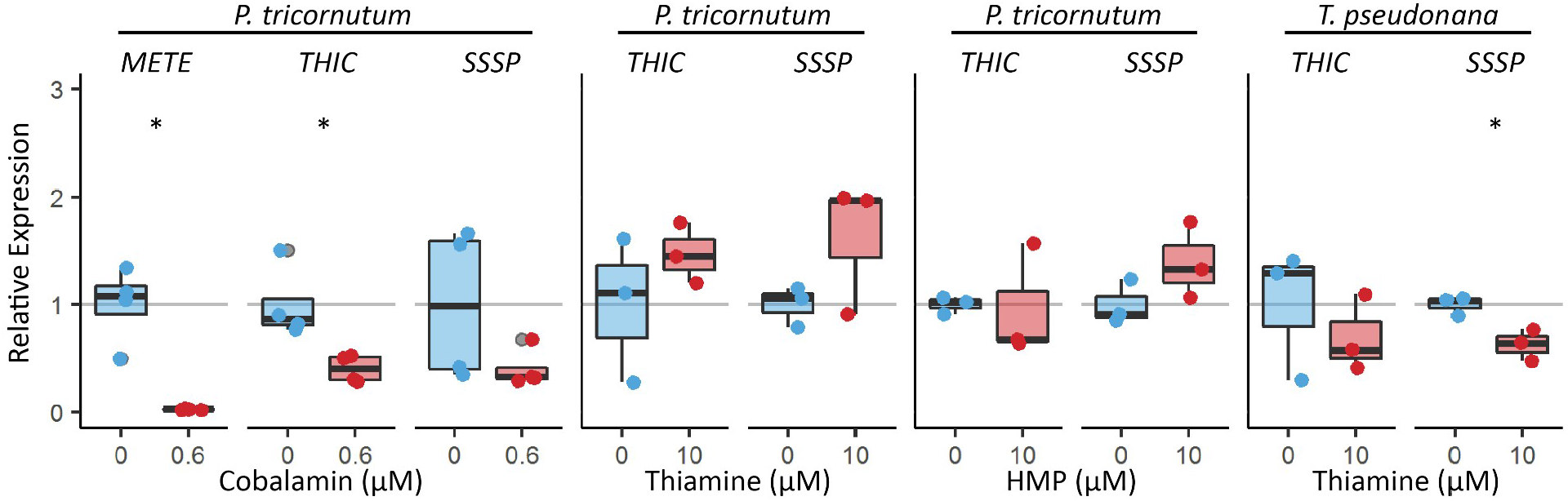
Impact of vitamin supplementation on expression of THIC and SSSP in Phaeodactylum tricornutum and Thalassiosira pseudonana. P. tricornutum and T. pseudonana were grown in the absence (blue) or presence (red) of 0.6 μM cobalamin (B_12_), 10 μM thiamine (B_1_) or 10 μM 4-Amino-5-hydroxymethyl-2-methylpyrimidine (HMP) for 7 days. Three or four biological replicates were analysed by RT-qPCR in technical duplicate. The technical replicate measurements were averaged for each biological replicate, and transcript levels were normalised for the average transcript levels of three housekeeping genes (H4, UBC, UBQ for P. tricornutum; Actin, EF1a, rbcs for T. psuedonana). Each dot represents the relative expression value for an individual biological replicate and a box plot summarises the data for each gene and treatment. Two-sided t-tests between supplemented and control conditions were conducted for all genes. *p-value < 0.05.

Acknowledging that gene regulation could also happen post-transcriptionally and given the presence of a predicted polyadenylation site overlapping the P1 stem in *PtTHIC*, we used a 3’RACE experiment to test whether the putative *PtTHIC* aptamer could regulate gene expression via alternative polyadenylation or alternative splicing. The results showed no substantive difference in *PtTHIC* 3’UTR isoforms between the control and the thiamine or HMP-supplemented conditions (Fig. **S5**). These results suggest that the *PtTHIC* predicted riboswitch does not regulate expression at a transcriptional or post-transcriptional level in response to thiamine.

Finally, to experimentally test whether the putative diatom aptamers can regulate thiamine metabolism, we employed a pyrithiamine growth assay, previously used to study thiamine gene regulation in other organisms (Sudarsan *et al*., 2005). Briefly, pyrithiamine, a thiamine antimetabolite, binds to the TPP aptamer downregulating the expression of thiamine biosynthesis genes regulated by TPP riboswitches, preventing the production of thiamine and inducing growth arrest. The lethal effect of pyrithiamine can be reversed by adding extracellular thiamine to compensate for the lack of biosynthetic activity. In this study, *C. reinhardtii, P. tricornutum* and *T. pseudonana* were grown in the presence or absence of 10 μM pyrithiamine and/or 10 μM thiamine (Fig. **4**). As can be seen clearly, *C. reinhardtii* growth is disrupted by pyrithiamine and rescued by thiamine supplementation, but *P. tricornutum* and *T. pseudonana* are insensitive to the antimetabolite.

**Figure 4.**
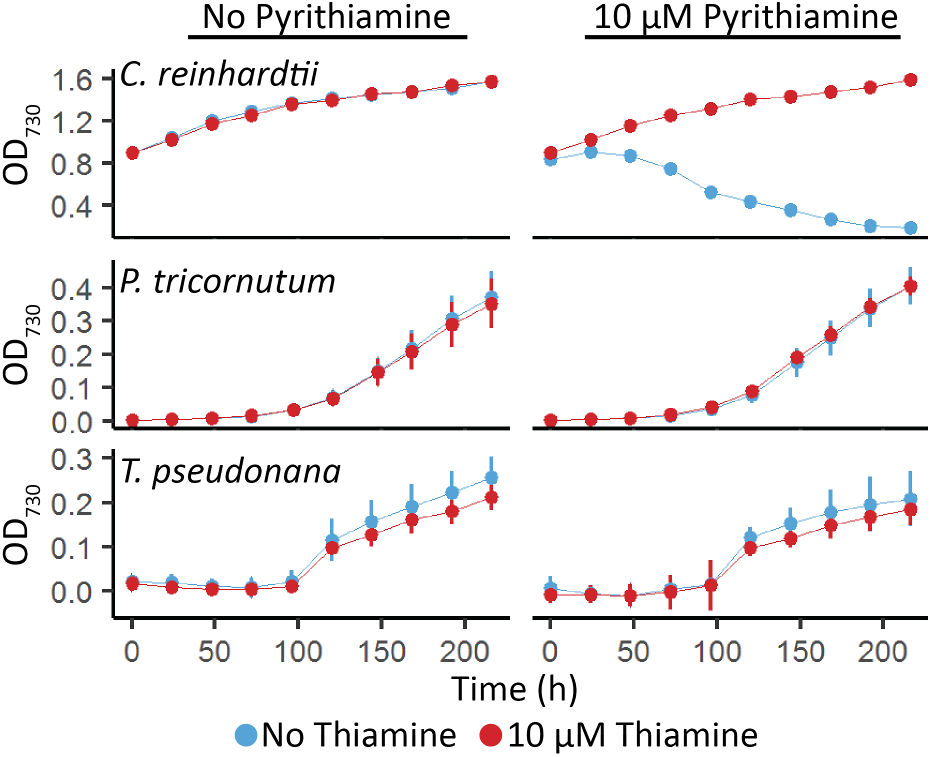
Effect of the thiamine antimetabolite pyrithiamine on the growth of Chlamydomonas reinhardtii, Phaeodactylum tricornutum and Thalassiosira pseudonana. C. reinhardtii, P. tricornutum and T. pseudonana were grown for 9 days in the absence (left column) or presence (right column) of 10 μM pyrithiamine and the absence (blue) or presence (red) of 10 μM thiamine in 96 well plates. Growth was measured as OD_730_ every 24 hours. Error bars represent the standard deviation of three biological replicates.

### The *P. tricornutum THIC* 3’UTR cannot regulate the expression of reporter constructs

As an alternative approach to determine whether the putative TPP aptamers in diatoms can regulate expression in response to thiamine supplementation, we generated and utilised a set of constructs where the putative *PtTHIC* riboswitch would regulate the expression of a reporter gene. We cloned the *PtTHIC* promoter, 5’UTR and 3’UTR so that they flanked a *Ble-Venus* reporter gene that confers resistance to zeocin. In principle, if the putative *PtTHIC* riboswitch regulated gene expression, the combined supplementation of thiamine and zeocin would induce a downregulation of the antibiotic-resistance reporter gene, which would in turn lead to growth arrest. However, we could not see any impact on growth when the transformants were cultured in the presence of 10 μM thiamine and 75 mg L^−1^ zeocin, providing further evidence that the putative *PtTHIC* riboswitch does not respond to thiamine supplementation (Fig. **5**).

**Figure 5.**
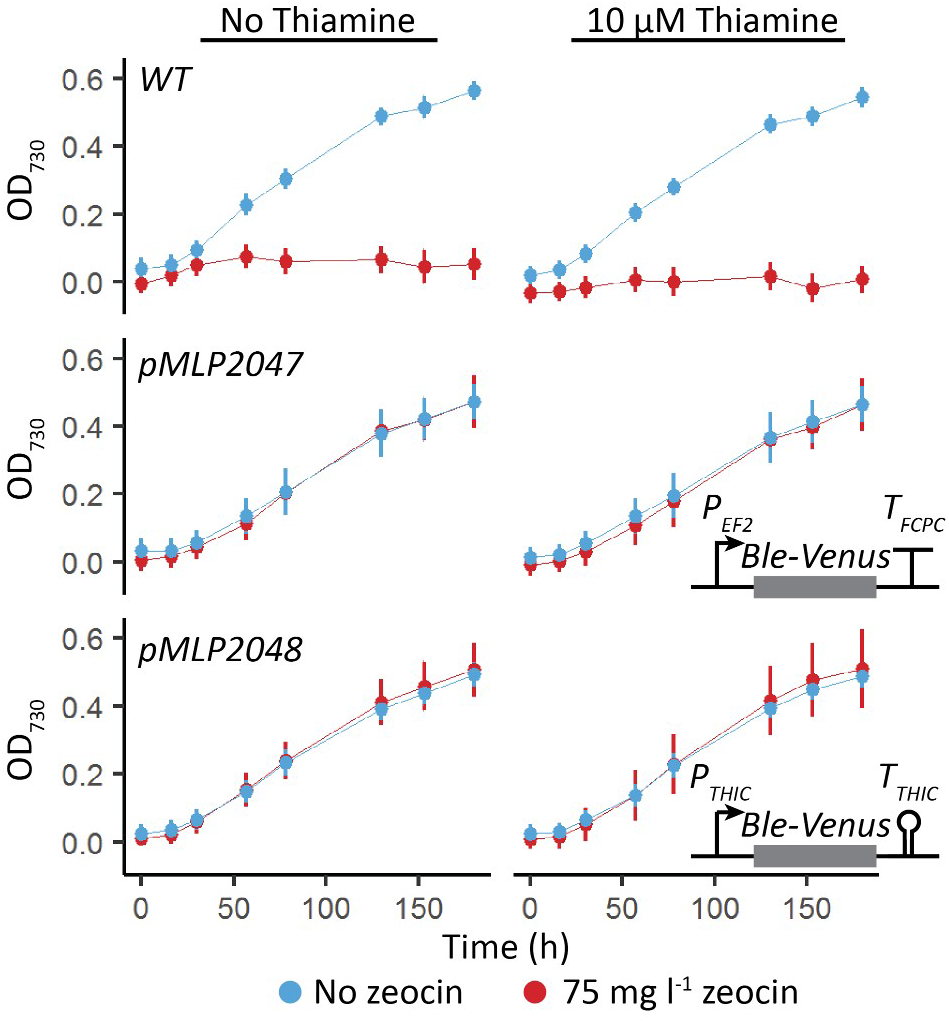
Effect of thiamine supplementation on transformants with PtTHIC promoter and 3’UTR driving expression of the Ble zeocin resistance gene. Transformants carrying a Ble-Venus reporter controlled by the PtEF2 promoter and PtFCPC 3’UTR (pMLP2047) or the PtTHIC promoter and 3’UTR (pMLP2048) were grown in the absence (left column) or presence (right column) of 10 μM and thiamine the absence (blue) or presence (red) of 75 mg L^−1^ zeocin. Error bars represent the standard deviation of three biological replicates for each of ten independent transformants for pMLP2047 and pMLP2048 and three biological replicates for WT.

*P. tricornutum* is unlikely to encounter thiamine concentrations at the micromolar level in oceanic environments where thiamine concentrations have been measured in the picomolar range (Sañudo-Wilhelmy *et al*., 2012; Monteverde *et al*., 2015). Thus, we wanted to test whether, despite being unresponsive to high levels of exogenous thiamine, the putative *PtTHIC* riboswitch is responsible for the homeostasis of intracellular TPP concentrations. To address this question, we employed a mutational approach inspired by previous observations in *A. thaliana* and *C. reinhardtii*, where mutations affecting functional TPP riboswitches in thiamine biosynthetic genes led to the overaccumulation of thiamine and TPP in response to a disruption of the negative feedback regulatory mechanism (Bocobza *et al*., 2013; Moulin *et al*., 2013). To replicate these experiments in *P. tricornutum*, we transformed WT cells with an extra copy of *PtTHIC* with a targeted mutation in the universally conserved pyrimidine-binding motif of the putative aptamer (“CUGAGA” to “CUCUCU”). To generate a control strain, a construct without this mutation was also transformed (Fig. **6a**). We then grew the transformants alongside a WT strain in the absence of exogenous thiamine for 5 days and quantified their intracellular thiamine and TPP levels by HPLC. The strains with the mutated copy of *PtTHIC* did not show any significant increase in intracellular thiamine or TPP compared to the unmutated control suggesting the putative *PtTHIC* aptamer is not required to regulate the homeostasis of thiamine and TPP levels (Fig. **6b**). In addition, the heterologous copies of *PtTHIC* in both constructs were tagged with a C-terminal HA-Tag so that we could follow changes in protein levels. If the riboswitch were functional, one would expect that a mutation in the universally conserved “CUGAGA” motif would disrupt feedback regulation and lead to increased protein levels. Western blot assays showed no visible increase in heterologous PtTHIC protein levels between the mutated and control constructs (Fig. **6c**). Furthermore, we saw no obvious changes in PtTHIC levels when 10 μM thiamine was added to transformants for the mutated or control constructs.

**Figure 6.**
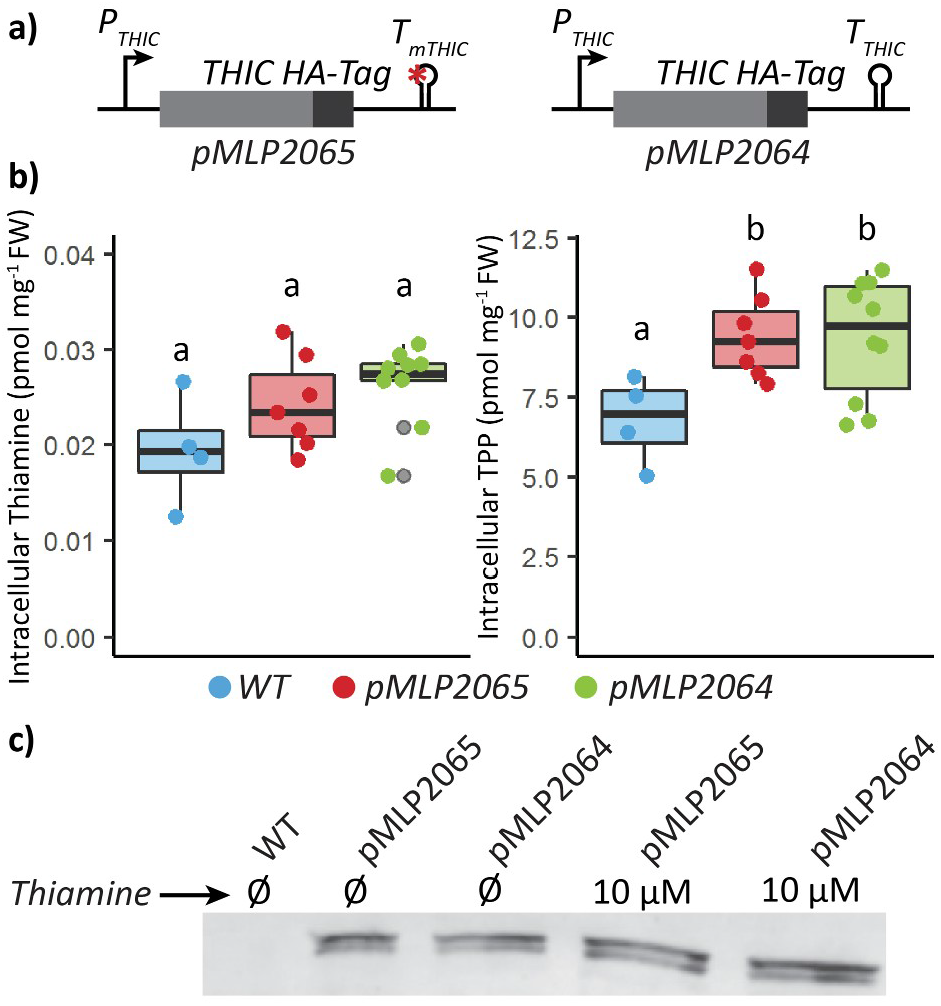
Intracellular thiamine and thiamine pyrophosphate (TPP) abundance and PtTHIC protein levels determined in transformants carrying a mutated PtTHIC aptamer. **(a)** A construct coding for an extra copy of PtTHIC with a targeted mutation in its putative aptamer (pMLP2065) and its respective unmutated control (pMLP2064) were transformed into Phaeodactylum tricornutum. **(b)** Transformants were grown for 5 days before thiamine and TPP were quantified by HPLC and normalised by fresh weight. Each dot represents the measurement of an independent transformant and a box plot summarises the data. Different letters represent significant differences in average vitamin content between strains in a Tukey HSD test with a 0.95 confidence level. **(c)** An independent transformant for each construct was grown to approximately 5×10^6^ cells mL^−1^ in the presence or absence of 10 μM thiamine, and protein was extracted from 150 mL cultures. A western blot analysis with a primary anti-HA antibody on total crude extracts normalised to culture OD is shown.

### The putative PtTHIC aptamer does not mediate switching in the CrTHI4 5’UTR aptamer platform

To test whether the putative *PtTHIC* aptamer was able to bind TPP and thereby regulate gene expression, we employed an aptamer testing platform that we recently developed in *C. reinhardtii* (Mehrshahi *et al*., 2020), which provides a simple measurable growth readout. Briefly, the aptamer platform allows the introduction of heterologous aptamers into a modified *CrTHI4* 5’UTR containing the riboswitch cloned in front of a *Ble-eGFP* reporter (Fig. **7a**). As before, if the introduced aptamers are functional in the platform context, the simultaneous presence of thiamine and zeocin in the medium impairs growth. In this study, we introduced the putative *PtTHIC* aptamer into the aptamer platform and employed the *CrTHIC* aptamer as a positive control to test whether the putative *PtTHIC* aptamer could respond to thiamine. We found that, as seen previously, the transformants with the *CrTHIC* aptamer showed impaired growth in the presence of thiamine and zeocin, with over a 3-fold difference in OD_730_ between the thiamine deplete and supplemented conditions four days post-inoculation (Fig. **7b**). In contrast, the transformants with the putative *PtTHIC* aptamer showed no growth difference between thiamine replete and deplete treatments. We then prepared a suite of modified aptamers combining functional domains from *CrTHIC* and *PtTHIC* aptamers to test whether a particular functional domain of the *PtTHIC* aptamer was responsible for the lack of thiamine response or was not compatible with the aptamer testing platform (Fig. **7a**). We found that neither introducing the P1 and P2 stems and/or the P4/5 stem from *CrTHIC* aptamer into the *PtTHIC* aptamer nor removing the P3a stem led to a responsive aptamer. In one of the modified aptamers, the only difference from the *CrTHIC* positive control was the L2/4 loop and the P3 stem from *PtTHIC* aptamer and yet this variant still failed to respond to thiamine (Fig. **7b**).

**Figure 7.**
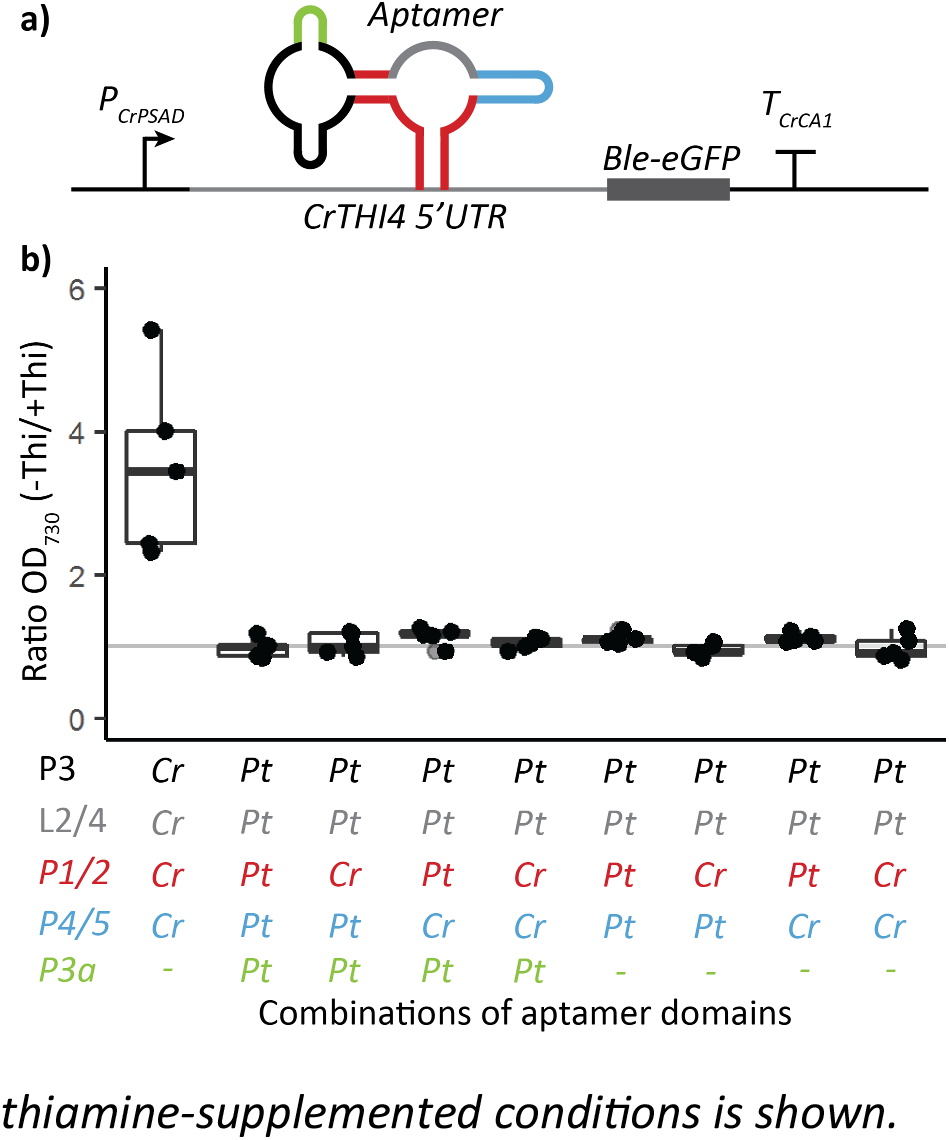
Response of Chlamydomonas reinhardtii carrying the PtTHIC-CrTHIC chimeric riboswitches to thiamine supplementation. **(a)** The CrTHI4 riboswitch platform previously developed in our lab (Mehrshahi et al., 2020) was cloned in the 5’UTR of a Ble-eGFP zeocin resistance reporter with a CrPSAD promoter and a CrCA1 terminator. A set of modified aptamers combining five structural parts (P1/2, P3, P3a, L2/4 and P4/5; colour coded) from CrTHIC and PtTHIC aptamers was cloned in the platform and the constructs transformed into C. reinhardtii. **(b)** Five independent transformants for each modified aptamer design were grown in the presence of 10 mg L^−1^ zeocin with or without 10 μM thiamine for four days. The ratio between the OD_730_ in the deplete and the thiamine-supplemented conditions is shown.

### *P. tricornutum SSSP* is necessary for thiamine uptake and *THIC* is essential for thiamine biosynthesis

Having observed that *PtTHIC* is not regulated by thiamine, unlike its homologues in bacteria, plants and chlorophytes, we sought to investigate whether the two *P. tricornutum* genes with putative TPP aptamers, *PtSSSP* and *PtTHIC*, were genuinely involved in thiamine metabolism in this diatom. We used CRISPR/Cas9-induced homologous directed repair to knock-out the genes and study their function. We generated knock-out strains for *PtTHIC* and *PtSSSP* by co-electroporating WT cells with a plasmid coding for Cas9 and a guide RNA pair, and another plasmid encoding a homologous repair template designed to swap the coding sequence of each target gene for a nourseothricin resistance cassette. This design facilitates genotypic screening by PCR and phenotypic screening by nourseothricin resistance (Fig. **8a,d**). After an initial screen of several hundred nourseothricin-resistant transformants, we identified several with insertions in the PtSSSP gene. Two were characterised further. Genotyping confirmed one mutant, called *ΔSSSP#1*, with a monoallelic disruption of the coding sequence around the sgRNA target sites, and a second mutant (*ΔSSSP#2*) with a biallelic disruption of the genomic sequence, in other words a complete knock-out (Fig. **8b**). When grown in the presence of 10 μM thiamine, intracellular thiamine levels in WT cells were substantially higher than in the absence of the vitamin (1.06 pmol mg^−1^ fresh weight (FW) compared to 0.13 pmol mg^−1^ FW), whereas in the *ΔSSSP#1* mutant the increase in intracellular thiamine was only to 0.72 pmol mg^−1^ FW (Fig. **8c**). There was no statistical difference in intracellular thiamine levels between *ΔSSSP#2* cells grown in the presence or absence of 10 μM thiamine (0.23 versus 0.09 pmol mg^−1^ FW) indicating that no exogenous thiamine had been taken up. This demonstrates that PtSSSP is essential for thiamine uptake and likely encodes a thiamine transporter (Fig. **8c**).

**Figure 8.**
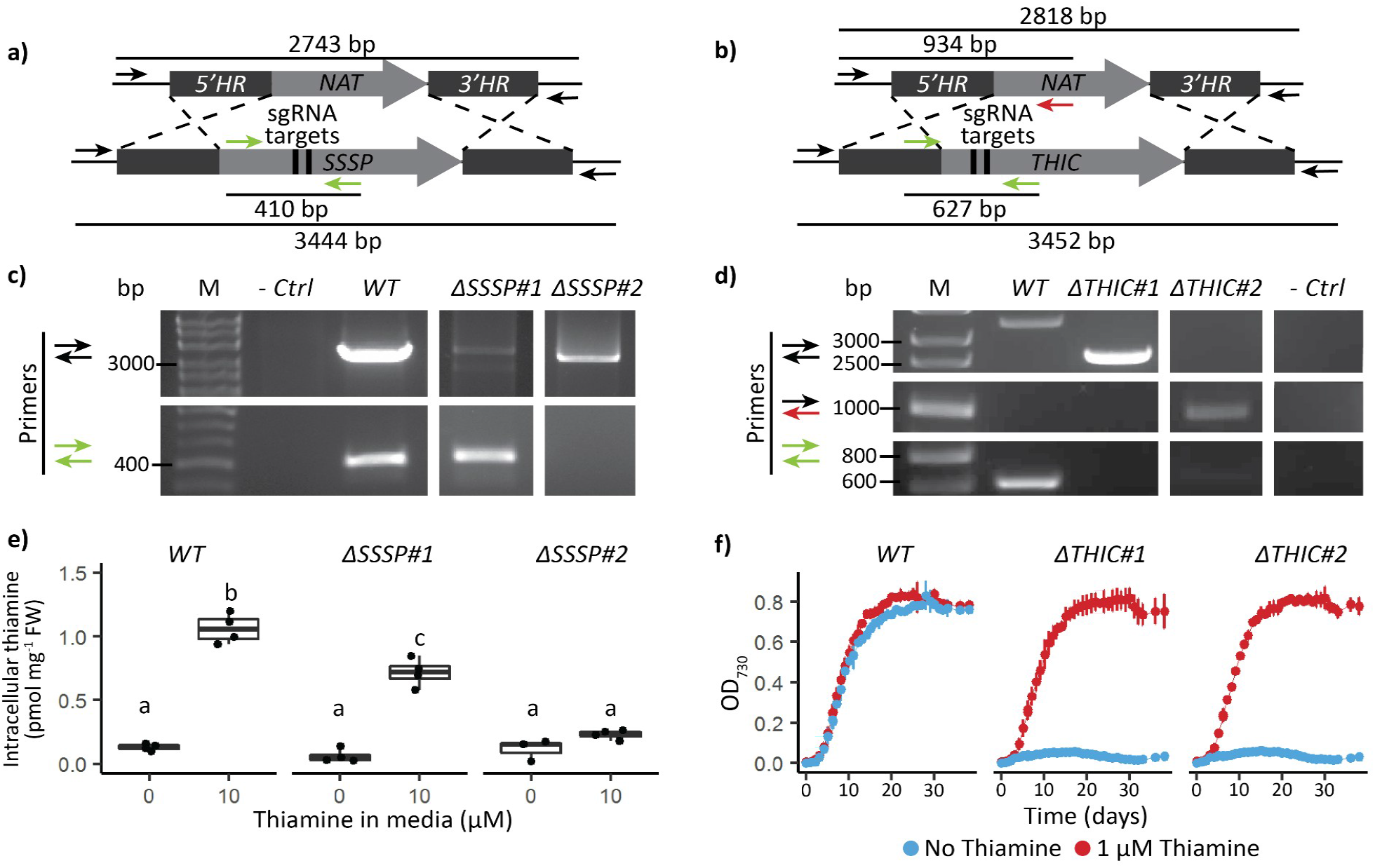
Determination of genotype and phenotype of Phaeodactylum tricornutum SSSP and THIC CRISPR/Cas9 mutants. **(a)** and **(b)** Schematic representation of the CRISPR-mediated homology recombination strategy to inactivate SSSP (Phatr3_J50012) and THIC (Phatr3_J38085), respectively. **(c)** and **(d)** Transformants were genotyped with two or three primer pairs colour-coded in panel a). The negative control did not include any template DNA. **(e)** WT and two SSSP knock-out strains were grown in the absence or presence of 10 μM thiamine for 5 days in biological duplicate, and intracellular thiamine levels were measured in technical duplicate. Different letters represent significant differences in average intracellular thiamine content between strains and conditions in a Tukey HSD test with a 0.95 confidence level. **(f)** WT and two THIC knock-out strains were grown in the absence (blue) or presence (red) of 1 μM thiamine in 24-well plates recording growth as OD_730_ every 24 hours. Error bars represent the standard deviation of three biological replicates.

For *THIC*, again after an initial screen of hundreds of transformants, we characterised two of them further. Both independent mutants showed a biallelic loss of the *PtTHIC* CDS (Fig. **8e**). While thiamine supplementation (at 1 μM) had no effect on growth of a WT control, both *ΔTHIC* mutants were able to grow only in the presence of thiamine, with no growth observed in its absence (Fig. **8f**). To confirm whether *PtTHIC* encodes an HMP-P synthase, we started three cultures of the *ΔTHIC#1* mutant in the absence of thiamine and at day 6 post-inoculation we supplemented the first culture with 1 μM thiamine, the second with 1 μM HMP, and the third was left unsupplemented. Both thiamine and HMP supplementation supported the growth of the mutant from that point (Fig. **9a**), confirming that *PtTHIC* encodes an HMP-P synthase. Finally, we grew the *ΔTHIC1* mutant in increasing concentrations of thiamine (0-500 nM) to establish the vitamin requirements of the mutant. As little as 5 nM thiamine was sufficient to support the growth of the mutant without detriment, but the mutant could not grow at 1 nM thiamine (Fig. **9b**).

**Figure 9.**
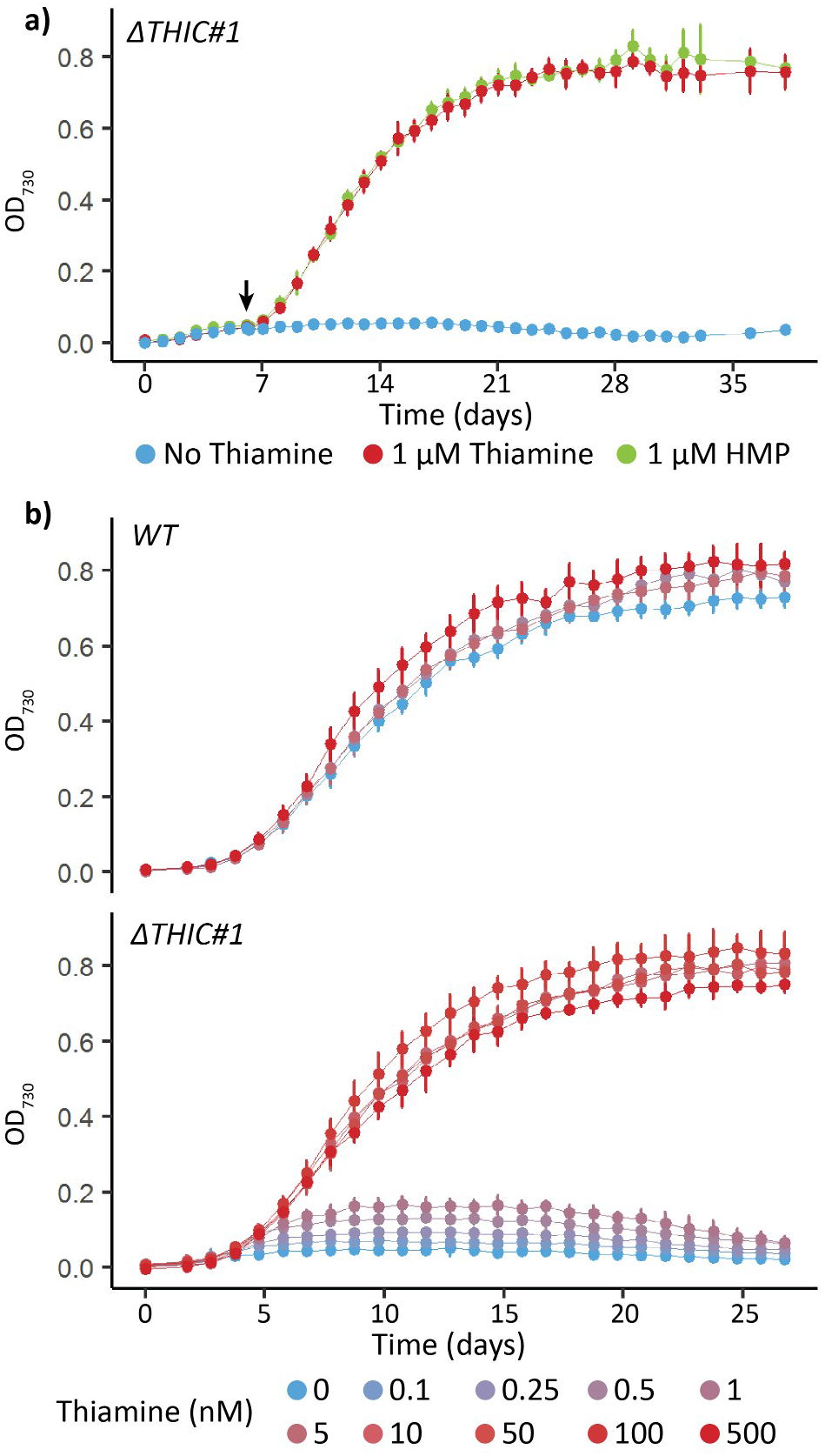
Phenotype analysis of a PtTHIC knock-out mutant. **(a)** The ΔTHIC#1 mutant was initially grown in the absence of supplementation, at day 6 (arrow) 1 μM thiamine (red) or 1 μM HMP was supplemented and growth compared to an unsupplemented control (blue). Error bars represent the standard deviation of three biological replicates. **(b)** WT and the ΔTHIC#1 mutant were grown in increasing concentrations of thiamine (0-500 nM) to determine the thiamine concentration required to support growth of the mutant. Error bars represent the standard deviation of six biological replicates.

## Discussion

Using a bioinformatics approach we screened over 20 published diatom genomes and found 41 previously unidentified putative TPP aptamer sequences, 32 of which were associated with four genes: *THIC, SSSP*, *SSUA/THI5-like* (encoding NMT1 domains), and *FOLR* (Fig. **2**, Table **S1b**). Riboswitches generally bind ligands related to the function of the genes they are physically associated with (McCown *et al*., 2017), suggesting that *THIC, SSSP*, *SSUA/THI5-like*, and *FOLR* are involved in thiamine metabolism. We have been able to validate experimentally the function of the first two genes in *P. tricornutum*, and yet the putative TPP aptamer sequences do not operate as riboswitches.

The thiamine auxotrophy shown by the *P. tricornutum THIC* knock-out mutants generated by CRISPR/Cas9 (Fig. **8f**) demonstrates that diatom *THICs* encode an HMP-synthase homologous to experimentally validated bacterial and plant HMP-synthases (Raschke *et al*., 2007; Chatterjee *et al*., 2008). Moreover, the inability of the *P. tricornutum ΔSSSP* mutant to uptake exogenous thiamine (Fig. **8c**) demonstrates, for the first time in an algal species, that SSSP is a thiamine transporter. SSSP belongs to the sodium-solute transporter (SSS) family, often associated with B-vitamin transporters (Jaehme & Slotboom, 2015). Members of this family have been found associated with predicted TPP riboswitches in chlorophytes, prasinophytes, cryptophytes, stramenopiles and haptophytes but, to our knowledge, have never been experimentally validated before (McRose *et al*., 2014). SSSP is homologous to the experimentally confirmed thiamine transporter Dur31 in the fungi *Candida parapsilosis*, which is also associated with a TPP aptamer (Donovan *et al*., 2018).

We also identified candidate *SSUA/THI5-like* genes containing NMT1 domains with homologies to the ThiY FAMP transporter (Bale *et al*., 2010) and the THI5 HMP-synthase (Coquille *et al*., 2012) in 9 diatom genomes. Except for the *F. solaris* candidates, all diatom candidates, together with those in haptophytes and cryptophytes, have a conserved signal peptide, which is present in ThiY but not THI5. In contrast, the multiple copies of the *F. solaris* candidate are phylogenetically more closely related to those in prasinophytes and chlorophytes, in agreement with their suggested horizontal gene transfer origin (Vancaester *et al*., 2020), and the fact that they are associated with predicted TPP aptamers. None of these have the signal peptide present in ThiY and SSUA, and therefore it is unlikely that they have the same function. Since the *P. tricornutum ΔTHIC* mutants cannot grow in the absence of thiamine (Fig. **8f**), the NMT1-domain containing gene is not sufficient for the production of HMP-P. Moreover, we observed that *P. tricornutum* CRISPR/Cas9 mutants lacking the NMT1 domain-containing gene are able to grow in the absence of exogenous thiamine and do not show an observable phenotype compared to WT, indicating that the NMT1-containing gene is not involved in the biosynthesis of thiamine (Fig. **S3**). In addition, the haptophyte *Emiliana huxleyi* has a NMT1 domain-containing gene but lacks *THIC* and is unable to grow without thiamine or the HMP derivative 4-amino-5-aminomethyl-2-methylpyrimidine (AmHMP; McRose *et al*., 2014; Gutowska *et al*., 2017). We propose that the previously used nomenclature *“SSUA/THI5-like”* (McRose *et al*., 2014) does not correspond to the actual function of NMT1-domain-containing genes in diatoms and should not be used. Instead we hypothesise that (with the exception of *F. solaris*) they are related to the bacterial *ThiY* and are involved in the salvage of the pyrimidine moiety. Further auxotrophy tests with NMT1 domain-containing gene mutants should be carried out to conclusively test whether these genes are involved in pyrimidine salvage. *FOLR* candidates were identified in 6 diatom genomes, and they are homologous to genes associated with predicted TPP aptamers in prasinophytes and rhizaria (McRose *et al*., 2014). Although the function of these genes remains unknown, they are predicted to have signal peptides, so they might be involved in thiamine transport through a receptor-mediated endocytosis mechanism similar to their homologous folate (vitamin B_9_) receptors in mice and humans (Zhao & Goldman, 2013).

The strong sequence conservation between putative diatom TPP aptamers, particularly in their P2, P4 and P5 stems and in the TPP binding motifs (Fig. 1a) indicate that the sequences have likely retained a defined and shared function within the diatom lineage. Given the absence of a conserved splicing acceptor site in P2 stems and the lack of evidence for alternative splicing in the *PtTHIC* 3’UTR in previous transcriptomic studies (Maheswari *et al*., 2009), our initial hypothesis was that the predicted diatom TPP riboswitches mechanism involved alternative polyadenylation in contrast to the alternative splicing mechanism shown for all previously characterised eukaryotic TPP riboswitches (Nguyen *et al*., 2016). This hypothesis was further supported by the conservation of A-rich P1 stems and the prediction of a polyadenylation site overlapping the *PtTHIC* aptamer P1 stem. In many eukaryotic genes, alternative polyadenylation determines differences in protein abundance, protein localisation, and/or protein-protein interactions between different transcript isoforms via the inclusion or exclusion of *cis*-regulatory elements bound by RNA binding proteins (Mayr, 2019).

However, despite the strong sequence conservation with experimentally characterised eukaryotic riboswitches and despite active thiamine uptake in *P. tricornutum* (Fig **S4**, Fig. **8c**), we were not able to demonstrate a change in transcript levels, nor alternative splicing or alternative polyadenylation in *PtTHIC* 3’UTR in response to thiamine supplementation (Fig. **3**, Fig. **S5**). The failure of *PtTHIC* 3’UTR to regulate a zeocin resistance reporter (Fig. **5**) and of the predicted aptamer to mediate a response to thiamine in the *CrTHI4* aptamer platform (Fig. **7**) support these observations and lead us to conclude that the predicted *PtTHIC* riboswitch does not regulate gene expression in response to thiamine supplementation. The stable levels of intracellular thiamine and TPP in transformants carrying a mutated *PtTHIC* aptamer (Fig. **6**) further demonstrate that in laboratory conditions the predicted *PtTHIC* riboswitch is not necessary to regulate the homeostasis of intracellular thiamine levels either. Although the qPCR results in *T. pseudonana* show a one-third downregulation of *TpSSSP* this change is only supported by a p-value of 0.03, and *TpTHIC* levels did not respond to thiamine (Fig. **3**). These qPCR results, together with the *P. tricornutum* and *T. pseudonana* resistance to pyrithiamine (Fig. **4**), suggest that the lack of response to thiamine supplementation by the predicted TPP riboswitches could be shared throughout the diatom lineage. It is worth noting that HMP can be obtained from the degradation of pyrithiamine (Sudarsan *et al*., 2005) and if the thiazol biosynthetic pathway is unaffected, the organisms would be able to survive despite THIC downregulation by pyrithiamine. We have not predicted any TPP riboswitches associated with the thiazol biosynthetic pathway in diatoms, hence the salvage of HMP could mask the results of the pyrithiamine experiment.

While all our experimental data coherently demonstrate that the predicted *PtTHIC* riboswitch does not respond to thiamine supplementation, the question remains why there is such sequence conservation across diatom aptamers, especially since these are in an untranslated region of the transcript. In general terms, we propose that while the riboswitch studied here may have lost the function of regulating gene expression in response to thiamine supplementation under laboratory conditions, it may have acquired a new functionality that has kept a high selection pressure. Taken together, our results show the weaknesses of bioinformatic approaches to predict riboswitch function and stress the necessity to experimentally test the functionality of the predicted aptamers before annotating them solely based on sequence or secondary structure conservation.

Thiamine is scarce in oceanic surface waters (Sañudo-Wilhelmy *et al*., 2012) and it is thought of as being growth-limiting for some primary producers in certain environments (Paerl *et al*., 2015), with special relevance for harmful algal species (Tang *et al*., 2010). In this context, the experimental confirmation of an HMP synthase and a thiamine transporter conserved in most of the available diatom genomes is of significant ecological relevance, given that these algae are responsible for 20% of global primary production (Field *et al*., 1998; Rousseaux & Gregg, 2014). The ubiquitous presence of genes encoding thiamine transporters and for the full thiamine biosynthesis pathway in the analysed diatom genomes does not offer sufficient information to hypothesise whether and in which conditions diatoms are net suppliers or consumers of thiamine and/or its moieties. The supply of thiamine and its moieties in oceanic environments has been shown to be dynamic and complex (Carini *et al*., 2014), and further research is needed to understand the ecological flows of this critical micronutrient in oceanic communities. Additionally, we have provided evidence to propose that genes encoding NMT1 domains found in several diatom, cryptophyte and haptophyte genomes are potentially involved in pyrimidine salvage. This is of special relevance given that some algal species have been shown to be dependent only on one of the thiamine moieties (Gutowska *et al*., 2017), and some marine bacterial species can grow on HMP but not on thiamine (Carini *et al*., 2014). Finally, we have found predicted TPP aptamers associated with most *THIC* and *SSSP* genes. Although our results show they are not responsive to thiamine supplementation under our laboratory conditions in *P. tricornutum* and *T. pseudonana*, we cannot rule out they have a conserved function significant for the regulation of thiamine metabolism with implications for thiamine dynamics in oceanic communities.

In summary, the findings presented here expand our knowledge on how thiamine is produced and taken up by diatoms and show the regulation of thiamine metabolism is more complex than previously thought. Further research will allow us to understand the full ecological and environmental implications of these findings in diatoms, a key taxonomic group in marine ecosystems and the main oceanic primary producers.

## Acknowledgments

We are grateful for technical support from Lorraine Archer. Jessie Dolliver helped to clone the HA-tagged copies of *THIC* and Astrid Stubbusch provided the zeocin selection cassette Level 1 construct used in most *P. tricornutum* constructs. Catherine Sutherland and A. Caroline Faessler helped to transform and maintain CRISPR/Cas9 mutants.

## Author Contributions

MLP and AGS conceived and designed the research; MLP, KG, AH, PM and AGS planned the experimental work; MLP, KG, PM, AH, SN and GIMO performed the experiments and data analysis; MLP and AGS wrote the manuscript with contributions from all authors. All authors reviewed and accepted the submitted manuscript.

## Data availability

All raw data, query sequences and scripts to generate the figures in this paper can be found in the GitHub online repository: https://github.com/AndreHolzer/Llavero-Pasquina_et_al_2021

## Supplementary Information

Additional Supplementary Information may be found online in the Supporting Information tab for this article:

**Fig. S1** Phylogenetic tree and multiple sequence alignment (MSA) for algal gene candidates with NMT1 domains.

**Fig. S2** Effect of thiamine and 4-Amino-5-hydroxymethyl-2-methylpyrimidine (HMP) on NMT1 domain-containing gene transcript levels in *Phaeodactylum tricornutum* and *Thalassiosira pseudonana*.

**Fig. S3** Characterisation of NMT1 domain-containing gene knock-out mutants generated by CRISPR/Cas9.

**Fig S4** *C. reinhardtii* and *P. tricornutum* intracellular thiamine and thiamine pyrophosphate (TPP) levels under increasing extracellular thiamine concentrations.

**Fig. S5** 3’RACE RT-PCR on *PtTHIC* in the presence or absence of 10 μM thiamine or 4-Amino-5-hydroxymethyl-2-methylpyrimidine (HMP).

**Table S1**. Thiamine pyrophosphate (TPP) riboswitch prediction in diatom genomes.

**Table S2**. Diatom genomes analysed in this study.

**Table S3**. Identification of thiamine-related genes in diatom genomes.

**Table S4**. Primers used in this study.

## References

Anthony PC, Perez CF, García-García C, Block SM. 2012. Folding energy landscape of the thiamine pyrophosphate riboswitch aptamer. Proceedings of the National Academy of Sciences, 109: 1485–1489.

Bale S, Rajashankar KR, Perry K, Begley TP, Ealick SE. 2010. HMP binding protein ThiY and HMP-P synthase THI5 are structural homologues. Biochemistry, 49: 8929–8936.

Beilharz TH, Preiss T. 2009. Transcriptome-wide measurement of mRNA polyadenylation state. Methods, 48: 294–300.

Bertrand EM, Allen AE. 2012. Influence of vitamin B auxotrophy on nitrogen metabolism in eukaryotic phytoplankton. Frontiers in microbiology, 3: 375.

Bertrand EM, Allen AE, Dupont CL, Norden-Krichmar TM, Bai J, Valas RE, Saito MA. 2012. Influence of cobalamin scarcity on diatom molecular physiology and identification of a cobalamin acquisition protein. Proceedings of the National Academy of Sciences, 109: E1762–E1771.

Bocobza SE, Malitsky S, Araújo WL, Nunes-Nesi A, Meir S, Shapira M, Fernie AR, Aharoni A. 2013. Orchestration of thiamin biosynthesis and central metabolism by combined action of the thiamin pyrophosphate riboswitch and the circadian clock in *Arabidopsis*. The Plant Cell, 25: 288–307.

Carini P, Campbell EO, Morré J, Sanudo-Wilhelmy SA, Thrash JC, Bennett SE, Temperton B, Begley T, Giovannoni SJ. 2014. Discovery of a SAR11 growth requirement for thiamin’s pyrimidine precursor and its distribution in the Sargasso Sea. The ISME journal, 8: 1727–1738.

Chatterjee A, Li Y, Zhang Y, Grove TL, Lee M, Krebs C, Booker SJ, Begley TP, Ealick SE. 2008. Reconstitution of ThiC in thiamine pyrimidine biosynthesis expands the radical SAM superfamily. Nature chemical biology, 4: 758–765.

Cheah MT, Wachter A, Sudarsan N, Breaker RR. 2007. Control of alternative RNA splicing and gene expression by eukaryotic riboswitches. Nature, 447: 497.

Coquille S, Roux C, Fitzpatrick TB, Thore S. 2012. The last piece in the vitamin b1 biosynthesis puzzle structural and functional insight into yeast 4-amino-5-hydroxymethyl-2-methylpyrimidine phosphate (hmp-p) synthase. Journal of Biological Chemistry, 287: 42333–42343.

Croft MT, Moulin M, Webb ME, Smith AG. 2007. Thiamine biosynthesis in algae is regulated by riboswitches. Proceedings of the National Academy of Sciences, 104: 20770–20775.

Croft MT, Warren, MJ, Smith AG. 2006. Algae need their vitamins. Eukaryotic cell, 5: 1175–1183.

Crozet P, Navarro FJ, Willmund F, Mehrshahi P, Bakowski K, Lauersen KJ, Pérez-Pérez ME, Auroy P, Gorchs Rovira A, Sauret-Gueto, S et al. 2018. Birth of a photosynthetic chassis: A MoClo toolkit enabling synthetic biology in the microalga Chlamydomonas reinhardtii. ACS synthetic biology, 7: 2074–2086.

Donovan PD, Holland LM, Lombardi L, Coughlan AY, Higgins DG, Wolfe KH, Butler G. 2018. TPP riboswitch-dependent regulation of an ancient thiamin transporter in *Candida*. PLoS genetics, 14: e1007429.

Eddy SR. 2011. Accelerated profile HMM searches. PLoS Comput Biol, 7: e1002195.

Engler C, Youles M, Gruetzner R, Ehnert TM, Werner S, Jones JD, Parton NJ, Marillonnet S. 2014. A golden gate modular cloning toolbox for plants. ACS synthetic biology, 3: 839–843.

Edgar RC. 2004. MUSCLE: multiple sequence alignment with high accuracy and high throughput, Nucleic Acids Research, 32: 1792–1797.

Field CB, Behrenfeld MJ, Randerson JT, Falkowski P. 1998. Primary production of the biosphere: integrating terrestrial and oceanic components. Science, 281: 237–240.

Gutowska MA, Shome B, Sudek S, McRose DL, Hamilton M, Giovannoni SJ, Begley TP, Worden AZ. 2017. Globally important haptophyte algae use exogenous pyrimidine compounds more efficiently than thiamin. MBio, 8: e01459–17.

Hanson AD, Amthor JS, Sun J, Niehaus TD, Gregory III JF, Bruner SD, Ding Y. 2018. Redesigning thiamin synthesis: Prospects and potential payoffs. Plant Science, 273: 92–99.

Hazra, AB, Han, Y, Chatterjee, A, Zhang, Y, Lai, RY, Ealick, SE, Begley, TP. 2011. A missing enzyme in thiamin thiazole biosynthesis: identification of TenI as a thiazole tautomerase. Journal of the American Chemical Society, 133: 9311–9319.

Helliwell KE, Wheeler GL, Smith AG. 2013. Widespread decay of vitamin-related pathways: coincidence or consequence?. Trends in Genetics, 29: 469–478.

Helliwell KE, Lawrence AD, Holzer A, Kudahl UJ, Sasso S, Kräutler B, Scanlan DJ, Warren MJ, Smith AG. 2016. Cyanobacteria and eukaryotic algae use different chemical variants of vitamin B12. Current Biology, 26: 999–1008.

Hofacker IL. 2003. Vienna RNA secondary structure server. Nucleic acids research, 31: 3429–3431.

Hopes A, Nekrasov V, Belshaw N, Grouneva I, Kamoun S, Mock T. 2017. Genome Editing in Diatoms Using CRISPR-Cas to Induce Precise Bi-allelic Deletions. Bio-protocol, 7: 23.

Hopes A, Nekrasov V, Kamoun S, Mock T. 2016. Editing of the urease gene by CRISPR-Cas in the diatom Thalassiosira pseudonana. Plant Methods, 12: 1–12.

Jaehme M, Slotboom DJ. 2015. Diversity of membrane transport proteins for vitamins in bacteria and archaea. Biochimica et Biophysica Acta (BBA)-General Subjects, 1850: 565–576.

Ji G, Li L, Li QQ, Wu X, Fu J, Chen G, Wu X. 2015. PASPA: a web server for mRNA poly (A) site predictions in plants and algae. Bioinformatics, 31: 1671–1673.

Jurgenson CT, Begley TP, Ealick SE. 2009. The structural and biochemical foundations of thiamin biosynthesis. Annual review of biochemistry, 78: 569–603.

Kanehisa M. 2019. Toward understanding the origin and evolution of cellular organisms. Protein Science, 28: 1947–1951.

Kanehisa M, Furumichi M, Sato Y, Ishiguro-Watanabe M, Tanabe M. 2021. KEGG: integrating viruses and cellular organisms. Nucleic Acids Research, 49: D545–D551.

Kanehisa M, Goto S. 2000. KEGG: kyoto encyclopedia of genes and genomes. Nucleic acids research, 28: 27–30.

Katoh K, Standley DM. 2013. MAFFT multiple sequence alignment software version 7: improvements in performance and usability. Molecular biology and evolution, 30: 772–780.

Kumar S, Stecher G, Li M, Knyaz C, Tamura K. 2018. MEGA X: Molecular Evolutionary Genetics Analysis across computing platforms. Molecular Biology and Evolution, 35: 1547–1549.

Maheswari U, Mock T, Armbrust EV, Bowler C. 2009. Update of the Diatom EST Database: a new tool for digital transcriptomics. Nucleic acids research, 37: D1001–D1005.

Mayr C. 2019. What are 3′ UTRs doing?. Cold Spring Harbor perspectives in biology, 11: a034728.

Maundrell K. 1990. nmt1 of fission yeast. A highly transcribed gene completely repressed by thiamine. Journal of Biological Chemistry, 265: 10857–10864.

McCown PJ, Corbino KA, Stav S, Sherlock ME, Breaker RR. 2017. Riboswitch diversity and distribution. Rna, 23: 995–1011.

McRose D, Guo J, Monier A, Sudek S, Wilken S, Yan S, Mock T, Archibald JM, Begley TP, Reyes-Prieto A, Worden AZ. 2014. Alternatives to vitamin B 1 uptake revealed with discovery of riboswitches in multiple marine eukaryotic lineages. The ISME journal, 8: 2517.

Mehrshahi P, Nguyen GTD, Gorchs Rovira A, Sayer A, Llavero-Pasquina M, Lim Huei Sin M, Medcalf EJ, Mendoza-Ochoa GI, Scaife MA, Smith AG. 2020. Development of novel Riboswitches for synthetic biology in the green Alga Chlamydomonas. ACS synthetic biology, 9: 1406–1417.

Mehta AP, Abdelwahed SH, Fenwick MK, Hazra AB, Taga ME, Zhang Y, Ealick SE, Begley TP. 2015. Anaerobic 5-hydroxybenzimidazole formation from aminoimidazole ribotide: An unanticipated intersection of thiamin and vitamin B12 biosynthesis. Journal of the American Chemical Society, 137: 10444–10447.

Monteverde DR, Gómez-Consarnau L, Cutter L, Chong L, Berelson W, Sañudo-Wilhelmy SA. 2015. Vitamin B1 in marine sediments: pore water concentration gradient drives benthic flux with potential biological implications. Frontiers in Microbiology, 6: 434.

Moulin M, Nguyen GT, Scaife MA, Smith AG, Fitzpatrick TB. 2013. Analysis of Chlamydomonas thiamin metabolism in vivo reveals riboswitch plasticity. Proceedings of the National Academy of Sciences, 110: 14622–14627.

Neupert J, Karcher D, Bock R. 2009. Generation of Chlamydomonas strains that efficiently express nuclear transgenes. The Plant Journal, 57: 1140–1150.

Nguyen GT, Scaife MA, Helliwell KE, Smith AG. 2016. Role of riboswitches in gene regulation and their potential for algal biotechnology. Journal of phycology, 52: 320–328.

Paerl RW, Bertrand EM, Allen AE, Palenik B, Azam F. 2015. Vitamin B1 ecophysiology of marine picoeukaryotic algae: strain-specific differences and a new role for bacteria in vitamin cycling. Limnology and Oceanography, 60: 215–228.

Palmer LD, Downs, DM. 2013. The thiamine biosynthetic enzyme ThiC catalyzes multiple turnovers and is inhibited by S-adenosylmethionine (AdoMet) metabolites. Journal of Biological Chemistry, 288: 30693–30699.

Petersen TN, Brunak S, Von Heijne G, Nielsen H. 2011. SignalP 4.0: discriminating signal peptides from transmembrane regions. Nature methods, 8: 785–786.

Raschke M, Bürkle L, Müller N, Nunes-Nesi A, Fernie AR, Arigoni D, Amrhein N, Fitzpatrick TB. 2007. Vitamin B1 biosynthesis in plants requires the essential iron–sulfur cluster protein, THIC. Proceedings of the National Academy of Sciences, 104: 19637–19642.

Rodionov DA, Vitreschak AG, Mironov AA, Gelfand MS. 2002. Comparative genomics of thiamin biosynthesis in procaryotes New genes and regulatory mechanisms. Journal of Biological chemistry, 277: 48949–48959.

Roth A, Breaker RR. 2009. The structural and functional diversity of metabolite-binding riboswitches. Annual review of biochemistry, 78: 305–334.

Rousseaux CS, Gregg WW. 2014. Interannual variation in phytoplankton primary production at a global scale. Remote sensing, 6: 1–19.

Sañudo-Wilhelmy SA, Cutter LS, Durazo R, Smail EA, Gómez-Consarnau L, Webb EA, Prokopenko MG, Berelson WM, Karl DM. 2012. Multiple B-vitamin depletion in large areas of the coastal ocean. Proceedings of the National Academy of Sciences, 109: 14041–14045.

Seppey M, Manni M, Zdobnov EM. 2019. BUSCO: assessing genome assembly and annotation completeness. In: Kollmar M, ed. Gene prediction. New York, NY: Humana Press, 227–245.

Sudarsan N, Cohen-Chalamish S, Nakamura S, Emilsson GM, Breaker RR. 2005. Thiamine pyrophosphate riboswitches are targets for the antimicrobial compound pyrithiamine. Chemistry & biology, 12: 1325–1335.

Tang YZ, Koch F, Gobler CJ. 2010. Most harmful algal bloom species are vitamin B1 and B12 auxotrophs. Proceedings of the national academy of sciences, 107: 20756–20761.

Vancaester E, Depuydt T, Osuna-Cruz CM, Vandepoele K. 2020. Comprehensive and functional analysis of horizontal gene transfer events in diatoms. Molecular Biology and Evolution, 37: 3243–3257.

Wachter A, Tunc-Ozdemir M, Grove BC, Green PJ, Shintani DK, Breaker RR. 2007. Riboswitch control of gene expression in plants by splicing and alternative 3′ end processing of mRNAs. The Plant Cell, 19: 3437–3450.

Webb ME, Marquet A, Mendel RR, Rébeillé F, Smith AG. 2007. Elucidating biosynthetic pathways for vitamins and cofactors. Natural product reports, 24: 988–1008.

White HB. 1976. Coenzymes as fossils of an earlier metabolic state. Journal of Molecular Evolution, 7: 101–104.

Winkler W, Nahvi A, Breaker RR. 2002. Thiamine derivatives bind messenger RNAs directly to regulate bacterial gene expression. Nature, 419: 95.

Yu Z, Geisler K, Leontidou T, Young RE, Vonlanthen, SE, Purton S, Abell C, Smith, AG. 2021. Droplet-based microfluidic screening and sorting of microalgal populations for strain engineering applications. Algal Research, 56: 102293.

Zhao R, Goldman ID. 2013. Folate and thiamine transporters mediated by facilitative carriers (SLC19A1-3 and SLC46A1) and folate receptors. Molecular aspects of medicine, 34: 373–385.

