## Supplementary Figures for "Thiamine metabolism genes in diatoms are not regulated by thiamine despite the presence of predicted riboswitches"

**Fig. S1** Phylogenetic tree and multiple sequence alignment (MSA) for algal gene candidates with NMT1 domains.

**Fig. S2** Effect of thiamine and 4-Amino-5-hydroxymethyl-2-methylpyrimidine (HMP) on NMT1 domain-containing gene transcript levels in *Phaeodactylum tricornutum* and *Thalassiosira pseudonana*.

**Fig. S3** Characterisation of NMT1 domain-containing gene knock-out mutants generated by CRISPR/Cas9.

**Fig S4** *C. reinhardtii* and *P. tricornutum* intracellular thiamine and thiamine pyrophosphate (TPP) levels under increasing extracellular thiamine concentrations.

**Fig. S5** 3'RACE RT-PCR on *PtTHIC* in the presence or absence of 10 µM thiamine or 4-Amino-5-hydroxymethyl-2-methylpyrimidine (HMP).

**Submitted separately:**

**Table S1.** Thiamine pyrophosphate (TPP) riboswitch prediction in diatom genomes.

**Table S2.** Diatom genomes analysed in this study.

**Table S3.** Identification of thiamine-related genes in diatom genomes.

**Table S4.** Primers used in this study.

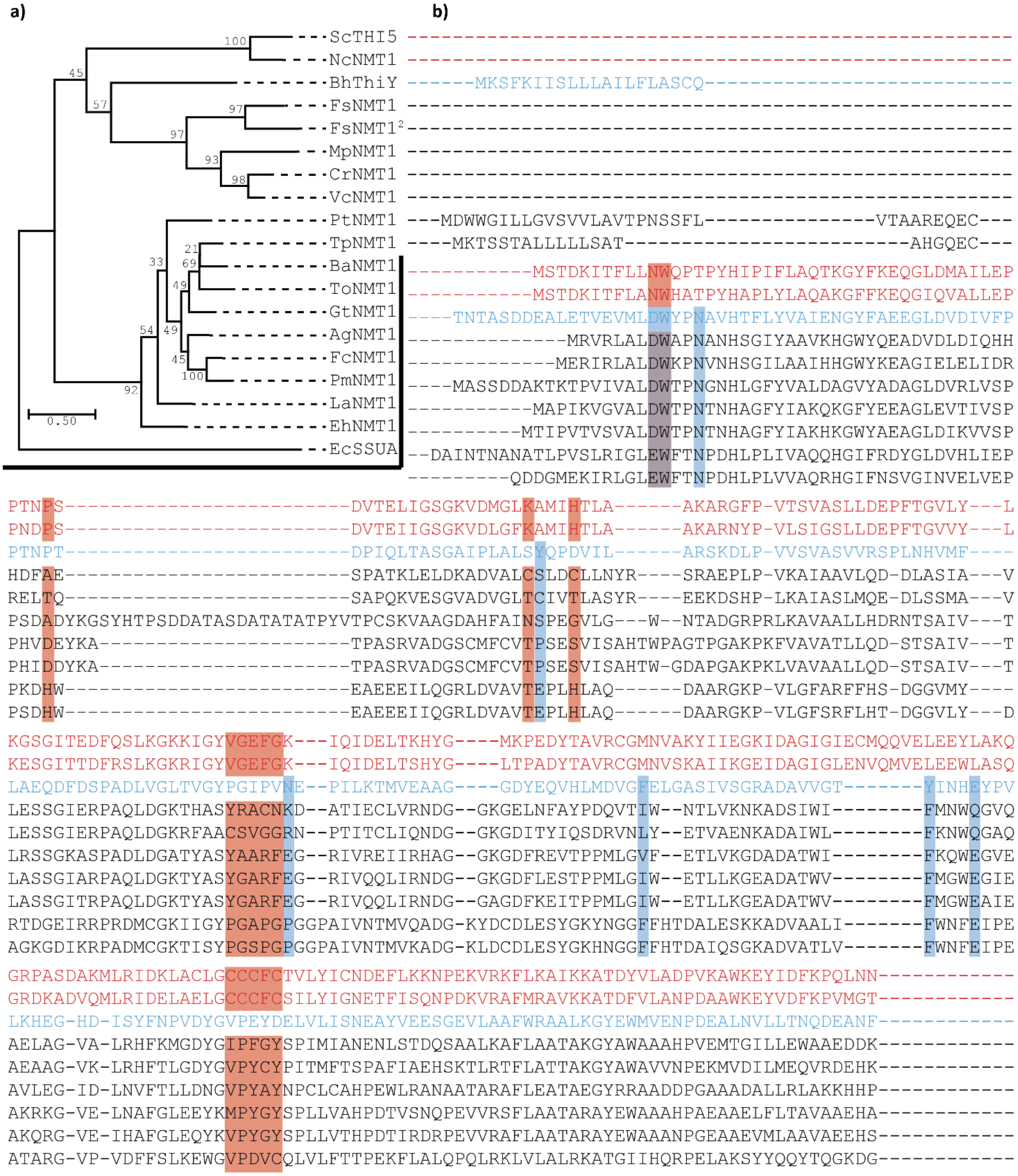


***Figure S1. Phylogenetic tree and multiple sequence alignment (MSA) for algal gene candidates with NMT1 domains. a)*** *The phylogenetic tree was generated with a Maximum-Likelihood algorithm and 100 bootstrap iterations (bootstrap values are indicated for each branch in the tree).* ***b)*** *The first 287 positions of the MSA are shown for the algal candidates with NMT1 domains. The alignment includes the bacterial thiamine transporter* BhThiY *(in blue letters), the fungal* NcNMT1 *and the yeast* ScTHI5 *homologues (in red letters). The peptide*

*sequences of the bottom eight algal sequences and* E. coli SSUA*, added as an outgroup for the tree, are not shown in the MSA. The alignment was generated in MEGA-X v.10.1.1 with MUSCLE. The active site residues of* BhThiY, *described in Bale* et al. *(2010), are highlighted in blue boxes, and the active site residues for* ScTHI5, *described in Coquille* et al. *(2012), are highlighted in red boxes to evaluate the conservation of key residues in the NMT1 domain-containing algal candidates. Sc*: Saccharomyces cerevisiae; *Nc*: Neurospora crassa; *Bh*: Bacillus halodurans; *Fs*: Fistulifera solaris *(two NMT1 domain containing peptides included, marked with a* [2]*)*; *Mp*: Micromonas pusilla; *Cr*: Chlamydomonas reinhardtii; *Vc*: Volvox carteri; *Pt*: Phaeodactylum tricornutum; *Tp*: Thalassiosira pseudonana; *Ba:* Bacillariophyta sp. (ASM1036717v1)*; To:* Thalassiosira oceanica*; Gt*: Guillardia theta; *Ag:* Asterionellopsis glacialis; *Fc:* Fragilariopsis cylindrus; *Pm:* Pseudo-nitzschia multiseries; *La:* Licmophora abbreviate; *Eh*: Emiliania huxleyi; *Ec*: Escherichia coli.

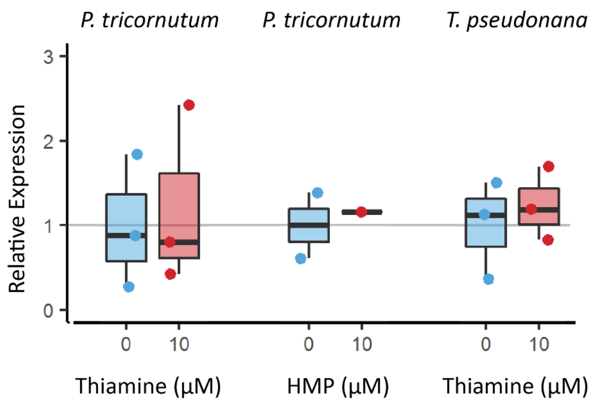


***Figure S2. Effect of thiamine and 4-Amino-5-hydroxymethyl-2-methylpyrimidine (HMP) on NMT1 domain-containing gene transcript levels in* Phaeodactylum tricornutum *and* Thalassiosira pseudonana.** *P. tricornutum and T. pseudonana were grown in the absence (blue) or presence (red) of 10 µM thiamine ($B_1$) or 10 µM 4-Amino-5-hydroxymethyl-2-methylpyrimidine (HMP) for 7 days. Three biological replicates were analysed by RT-qPCR in technical duplicate. The technical replicate measurements were averaged for each biological replicate, and transcript levels were normalised for the average transcript levels of three housekeeping genes (H4, UBC, UBQ for P. tricornutum; Actin, EF1a, rbcs for T. psuedonana). Each dot represents the relative expression value for an individual biological replicate and a box plot summarises the data for each gene and treatment. Two-sided t-tests between supplemented and control conditions were conducted for all genes. *p-value < 0.05.*

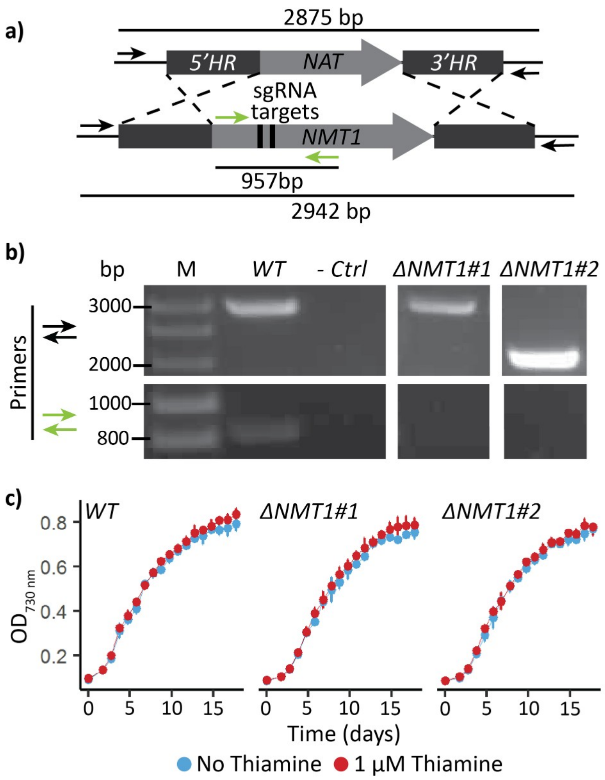


***Figure S3. Characterisation of NMT1 domain-containing gene knock-out mutants generated by CRISPR/Cas9*. a)** *Schematic representation of the CRISPR-mediated homology recombination strategy to inactivate the NMT1 domain-containing gene (*Phatr3_J33535*).* ***b)*** *Two mutants were genotyped alongside WT with two primer pairs colour-coded in panel a). The negative control did not include any template DNA* ***c)*** *Both mutants and the WT were cultured in 96-well plates in the absence (blue) or presence (red) of 1 µM thiamine in biological triplicate measuring $OD_{730nm}$ daily. Error bars represent the standard deviation of three biological replicates.*

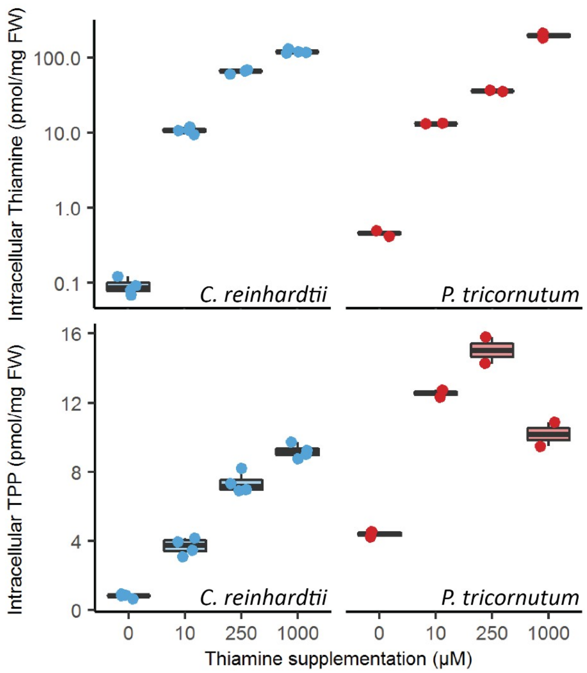


***Figure S4.*** **C. reinhardtii *and* P. tricornutum *intracellular thiamine and thiamine pyrophosphate (TPP) levels under increasing extracellular thiamine concentrations.*** *Cells were grown under increasing concentrations of extracellular thiamine for 5 days, then they were harvested, weighted and their intracellular thiamine and TPP levels determined by HPLC analysis. Two biological and two technical replicates for C. reinhardtii (blue dots) and two technical replicates were measured for P. Tricornutum (red dots).*

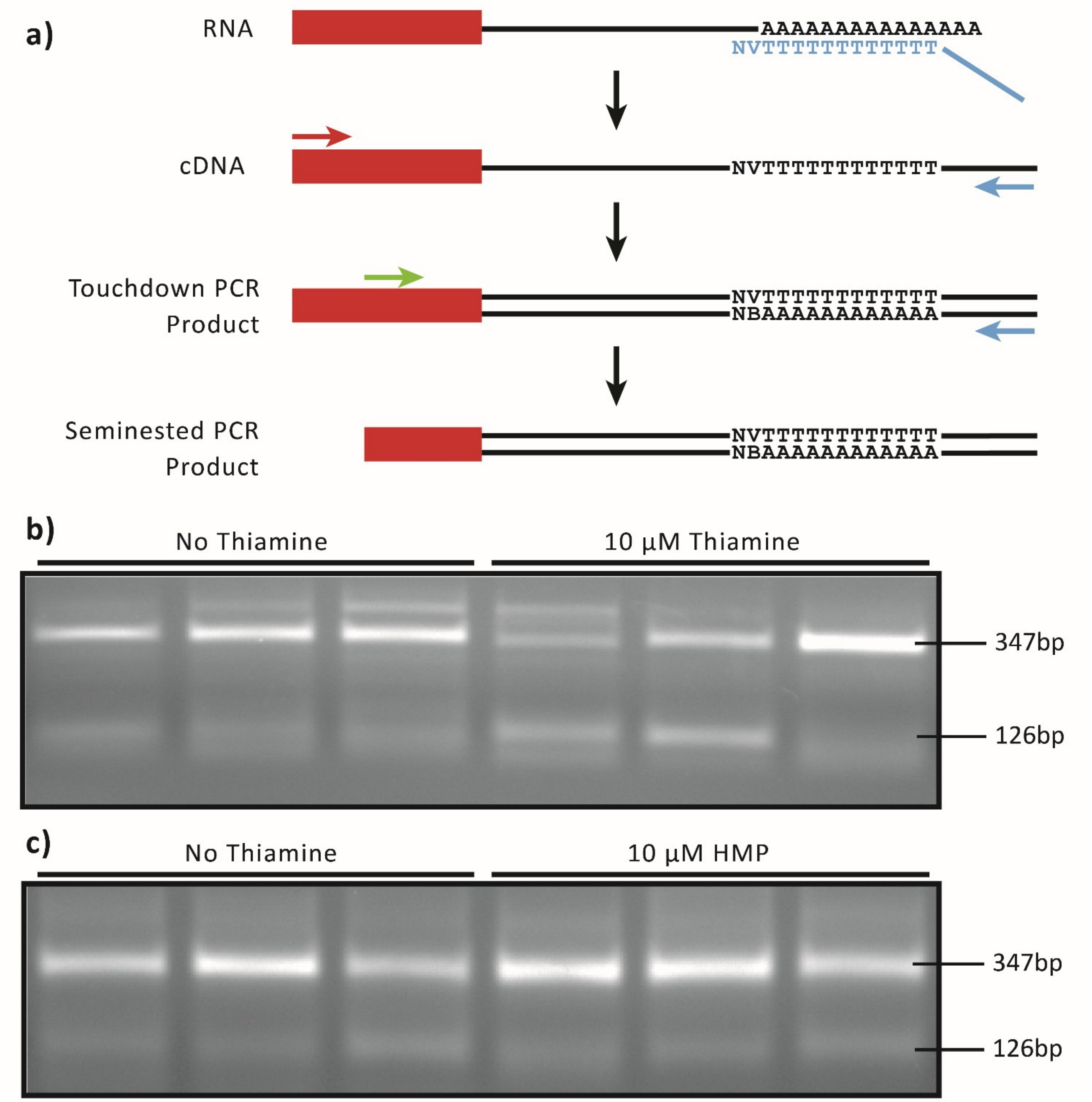


***Figure S5. 3'RACE RT-PCR on* PtTHIC *in the presence or absence of 10 µM thiamine or 4-Amino-5-hydroxymethyl-2-methylpyrimidine (HMP). a)*** *RNA isolated from P. tricornutum triplicate cultures in the presence or absence of 10 µM thiamine* ***(b)*** *or HMP* ***(c)*** *was retrotranscribed with polyT-VN primers binding the 5' end of the polyA site of messenger RNAs. A first touchdown PCR with a PtTHIC specific primer (red arrow) binding the coding sequence (red box) and a universal primer binding the adaptor included in the polyT-VN primer (blue arrow) was followed by a semi-nested PCR with a second PtTHIC specific primer (green arrow) and the universal primer to increase the specificity of the 3'RACE. Products of this second PCR*

*were run in a 0.5 % agarose gel. The length of the bands was established by Sanger sequencing. The 347 bp band indicates a polyadenylation site consistent with the available EST tags (Maheswari et al., 2019) and the polyadenylation site predicted by PASPA (Ji et al., 2015). The shorter band of 126 bp could be a technical artefact arising from the annealing of the TVN primer to an adenine-rich sequence in the 3'UTR (5' – AAAATAGA – 3').*

### References Supplementary Figures


Bale S, Rajashankar KR, Perry K, Begley TP, Ealick SE. 2010. HMP binding protein ThiY and HMP-P synthase THI5 are structural homologues. *Biochemistry*, *49:* 8929-8936.

Coquille, S., Roux, C., Fitzpatrick, T. B., & Thore, S. 2012. The last piece in the vitamin b1 biosynthesis puzzle structural and functional insight into yeast 4-amino-5-hydroxymethyl-2-methylpyrimidine phosphate (hmp-p) synthase. *Journal of Biological Chemistry*, *287:* 42333-42343.

Ji G, Li L, Li QQ, Wu X, Fu J, Chen G, Wu X. 2015. PASPA: a web server for mRNA poly (A) site predictions in plants and algae. *Bioinformatics*, *31:* 1671-1673.

Maheswari U, Mock T, Armbrust EV, Bowler C. 2009. Update of the Diatom EST Database: a new tool for digital transcriptomics. *Nucleic acids research*, *37:* D1001-D1005.
